# Spatial analysis reveals the evolving organization of low-grade and high-grade IDH-mutant glioma

**DOI:** 10.64898/2026.03.12.709536

**Authors:** Rouven Hoefflin, Alissa C. Greenwald, Noam Galili Darnell, Christopher W. Mount, Yali Tiomkin, Dor Simkin, Angelica B. Patterson, L. Nicolas Gonzalez Castro, Inna Goliand, Ofra Golani, Kevin Joseph, Jürgen Beck, Vidhya M. Ravi, Merav Kedmi, Hadas Keren-Shaul, Yoseph Addadi, Marian C. Neidert, Mario L. Suvà, Itay Tirosh

**Author notes:** Corresponding authors (I.T.), (M.L.S.). These authors contributed equally (Rouven Hoefflin, Alissa C. Greenwald, Noam Galili Darnell, Christopher W. Mount).

## Abstract

Adult diffuse gliomas are comprised of malignant cell states interwoven with the non-malignant brain microenvironment. Here we combine spatial transcriptomics and spatial proteomics of IDH–mutant gliomas to define organizational principles across histological grades. In low-grade tumors, spatial organization arises from underlying nonmalignant brain structures. For example, we classify low-grade tumor regions as embedded into white matter and identify a sharp white–grey matter junction that restricts cortical invasion and is associated with marked changes in tumor composition and cellular phenotypes. This junction is preferentially traversed by oligodendrocyte progenitor (OPC)-like malignant cells, which may drive tumor expansion. In contrast, intermediate-grade tumors are largely disorganized, with few recurring pairwise interactions between cancer cell states and TME cell types. In high-grade tumors, hypoxia/necrosis-associated global structure begins to emerge, reminiscent of IDH-wildtype glioblastoma. Together, these findings reveal two independent axes of glioma spatial organization—from brain anatomy-driven organization in low-grade tumors to hypoxia-associated structure in high-grade tumors—and establishes a framework that links tumor grade to recurrent spatial associations between cell states and cell types.

## Introduction

IDH–mutant (IDH-mut) gliomas frequently exhibit an initial period of indolent growth followed by inevitable recurrence, aggressive expansion, and treatment resistance, underscoring the need to understand the cellular and microenvironmental features that shape their progression^1^. The two major types of IDH-mut glioma—astrocytoma (IDH-A) and oligodendroglioma (IDH-O)—are distinguished by histological and genetic hallmarks, including loss of *ATRX* in IDH-A and chromosome 1p/19q co-deletion in IDH-O^2^. Despite these distinctions, single-cell RNA-sequencing (scRNA-seq) studies have shown that both tumor types share the same malignant cellular states and inferred hierarchy. By contrast, there is distinct variability in the composition of the non-malignant tumor microenvironment (TME) between the two entities^3,4^.

IDH-A and IDH-O are further classified into WHO grades based on integrated histological and genetic criteria^2^, which correlate with tumor aggressiveness and prognosis. Yet, despite the centrality of grading in clinical practice, our understanding of how malignant cell states and TME populations differ between grades remains incomplete^5,6^. Even less is known about how these cellular programs are spatially organized across tumor grades, or how spatial organization is generated and maintained during glioma evolution, as spatial profiling on IDH-mutant glioma thus far has focused on the immune microenvironment^7,8^.

Recent work in IDH-wildtype glioblastoma (GBM), the most common high-grade glioma of adults, has shown that hypoxia and necrosis are dominant environmental forces that drive spatial organization of GBM tissue architecture^9–11^. In these tumors, structured regions exhibit clusters of similar malignant and immune states that assemble into a recurring five-layered architecture in proximity to hypoxia/necrosis, whereas disorganized regions display a mixture of these cell states with fewer recurring spatial patterns.

In contrast, low-grade IDH-mut gliomas lack necrosis and might therefore be expected to display minimal spatial structure. Moreover, it is unknown whether—and to what extent—the highly organized architecture of the normal brain, including the distinct histology of grey and white matter, influences spatial patterns within these infiltrative tumors.

To address these gaps, we performed multimodal spatial profiling of IDH-mut gliomas across WHO CNS 5 grades using 10x Visium spatial transcriptomics and CODEX spatial proteomics and integrated these datasets with our previously published spatial GBM study^9^. By quantitatively assessing organization and identifying recurring spatial associations among malignant and non-malignant cellular states, we reveal how the patterns and drivers of spatial organization shift across glioma grades. Finally, we uncover an unexpected functional barrier in the white–grey matter junction in lower-grade IDH-mut gliomas. This anatomical interface is characterized by an abrupt change in TME cells and cancer cell states and may be preferentially traversed by a distinct malignant OPC-like cell state.

## Results

### Spatial proteomics and transcriptomics of IDH-mut glioma

We performed spatial proteomics and transcriptomics on a cohort of 34 primary IDH-mut glioma samples from 28 patients. Specifically, we applied CODEX to 30 samples, comprising approximately 2.6 million cells, and 10x Visium to 24 samples, with 63,839 spots (**Fig. 1A, Table S1**). In most cases (n = 20 samples), adjacent tissue sections were profiled by both technologies (**Fig. 1B**). Our cohort included the two IDH-mut glioma types - astrocytoma (IDH-A, n = 23) and oligodendroglioma (IDH-O, n = 11) - spanning WHO grades 2, 3, and 4 (**Fig. 1B**). These grades combine morphological and molecular features (e.g., *CDKN2A* loss)^2^ that correlate with prognosis and provide a unique framework to study spatial grade characteristics in glioma. To jointly analyze the spatial architecture of IDH-A and IDH-O in the available samples, we established a *categorical grade* based on histological features following the WHO-CNS5 guidelines: *low-grade* for grade 2 IDH-mut gliomas, *high-grade* for IDH-mut gliomas with evidence of necrosis and/or microvascular proliferation (MVP), and *intermediate-grade* (*mid-grade*) for all other IDH-mut gliomas. IDH-wt GBM (by definition grade 4) was analyzed as a separate category and was incorporated from Ravi et al.^11^ and our previously published IDH-wt GBM dataset (26 Visium samples with 65,995 spots; 12 CODEX samples with 428,395 cells), enabling a comprehensive spatial comparison across the most common types of adult diffuse glioma (**Table S1**).

**Figure 1.**
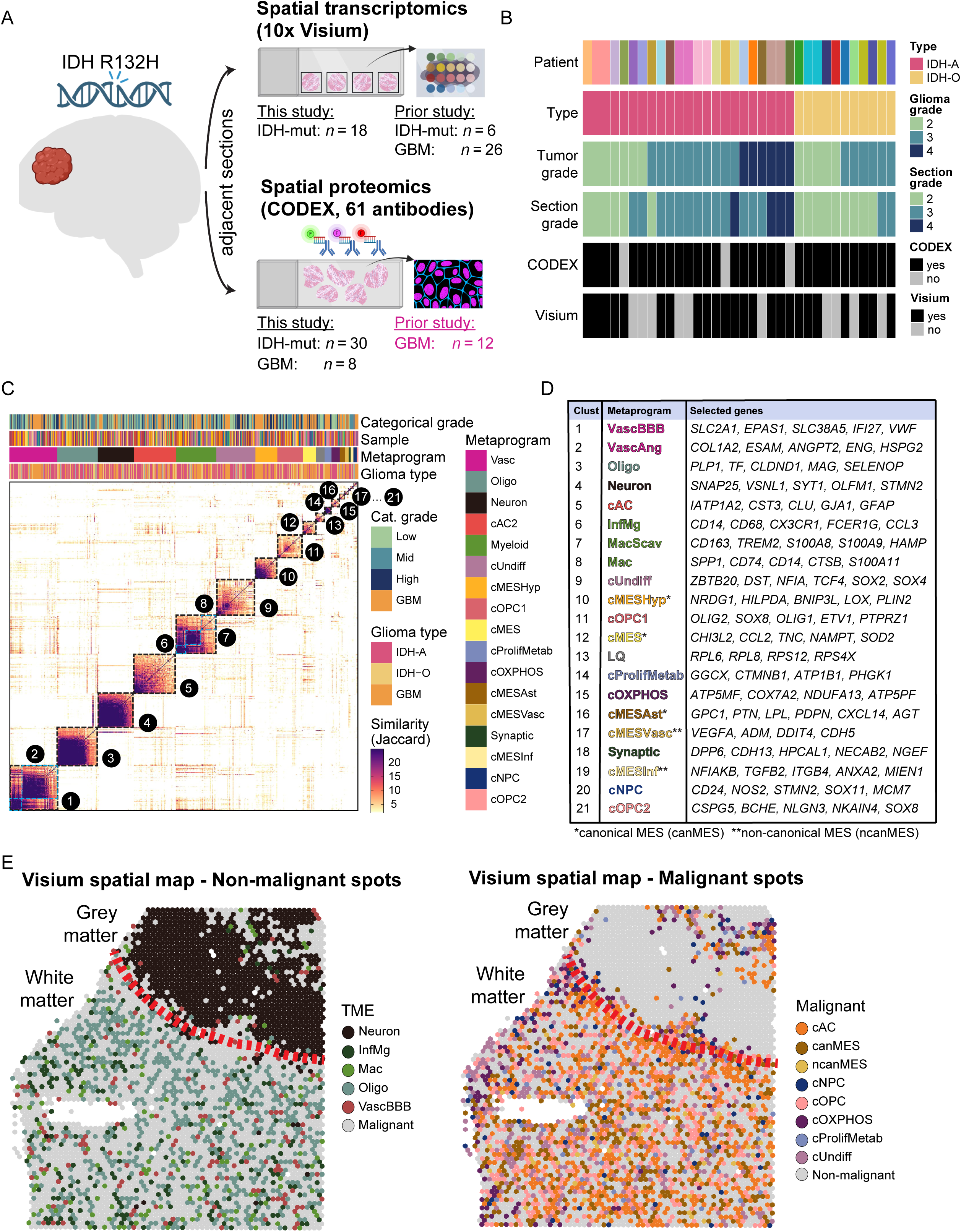
Experimental design and metaprogram generation. **(A)** Experimental overview and patient cohort. Fresh-frozen tissue sections from IDH-mut gliomas (n = 30) and IDH-wt GBM (n = 8) were profiled by CODEX using a 61-antibody panel. 10x Visium spatial transcriptomics was performed on adjacent sister sections (n = 24). This dataset was integrated with previously published spatial datasets^9,11^. Schematic created with BioRender.com. **(B)** Summary of the patient cohort including standard neuropathological classification, WHO grade based on tumor, and WHO grade based on the analyzed section. **(C)** Similarity matrix based on gene-set overlap (Jaccard index) across all transcriptional programs derived from NMF and Leiden clustering of the combined IDH-wt and IDH-mut datasets. Programs are annotated by sample, glioma type, categorial grade, and MP; cluster numbers correspond to those in (D). **(D)** Table of MPs showing representative genes for each numbered cluster in (C). Malignant MPs are prefixed with “c”. **(E)** Visium spatial maps displaying the spatial distribution of MPs stratified by non-malignant TME and malignant MPs.

### Identifying glioma cell states and non-malignant cell types of the TME

To identify variable gene metaprograms (MPs) that reflect recurrent cell states and cell types found in glioma, we combined unsupervised clustering approaches - Leiden clustering and non-negative matrix factorization (NMF) - on the Visium data^9^. Two distinct sets of MPs were generated: one exclusively from IDH-mut samples (IDH-mutMP, **Fig. S1A,B**, **Table S2**), and the second by integrating IDH-mut and IDH-wt GBM samples (GliomaMP, **Fig. 1C,D, Table S2**), facilitating pan-cohort comparisons. We also defined a set of generalized MPs, by collapsing MPs with high gene expression overlap that corresponded to a single cell state or a single cell type (GeneralMP, **Fig. S1C**).

The GliomaMPs revealed the expected malignant cell states as previously defined by scRNA-seq of IDH-mut gliomas^3^, GBM^12^ and Visium^9^. Throughout the manuscript, malignant MPs are prefixed with “c” for cancer. These included programs corresponding to undifferentiated (cUndiff), astrocyte-like (cAC), oligodendrocyte progenitor-like (cOPC), neuronal progenitor-like (cNPC), and mesenchymal cell states (cMES) (**Fig. S1D**). Mesenchymal states further subdivided into those enriched with astrocytic genes (cMESAst), hypoxic genes (cMESHyp), or neither (cMES), and are collectively referred to as *canonical MES* (*canMES*). New malignant MPs uniquely emerging from IDH-mut gliomas were related to oxidative phosphorylation (cOXPHOS), and to mesenchymal programs enriched with vascular or inflammatory genes (cMESVasc, cMESInf, collectively referred to as *non-canonical MES, ncanMES*).

Non-malignant spatial MPs reflecting the tumor microenvironment (TME) included the cell types of the normal brain parenchyma, such as neurons (Neuron), oligodendrocytes (Oligo), and normal blood-brain-barrier related vasculature (VascBBB) (**Fig. S1E**). We also identified an aberrant vasculature program enriched for genes involved in angiogenesis and extracellular matrix deposition (VascAng), and three distinct myeloid MPs including Inflammatory-Microglia (InfMac), Tumor-associated Macrophages (Mac) and immunosuppressive scavenger macrophages (MacScav)^10^. Visium spots were annotated by the highest scoring MP, as previously described ^9^ (Methods, **Fig. 1E**, **Fig. S1F**).

As an orthogonal approach, we performed spatial proteomics using the co-detection by indexing (CODEX) platform (**Fig. 1A**) with a comprehensive antibody panel (n=61, **Table S3**) followed by conventional H&E staining. This allowed us to map glioma cells and non-malignant TME cells at true single-cell resolution (**Fig. 2A-C**, **Fig. S2A**) and in spatial context of local tumor burden using Ivy Glioblastoma Atlas Project (IvyGAP) anatomic regions as a guide (**Fig. S2B**). Cell identities were annotated through a combined supervised and unsupervised approach (Methods). Briefly, nuclear segmentation of multiplexed images was followed by scoring each cell for all markers using the recently developed deep-learning model Nimbus^13^. Nimbus scores of canonical cell type markers were first used to assign cells to major classes (e.g., malignant, T-cell, vascular, myeloid). Second, subclusters within each class were derived via unsupervised clustering (Methods).

**Figure 2.**
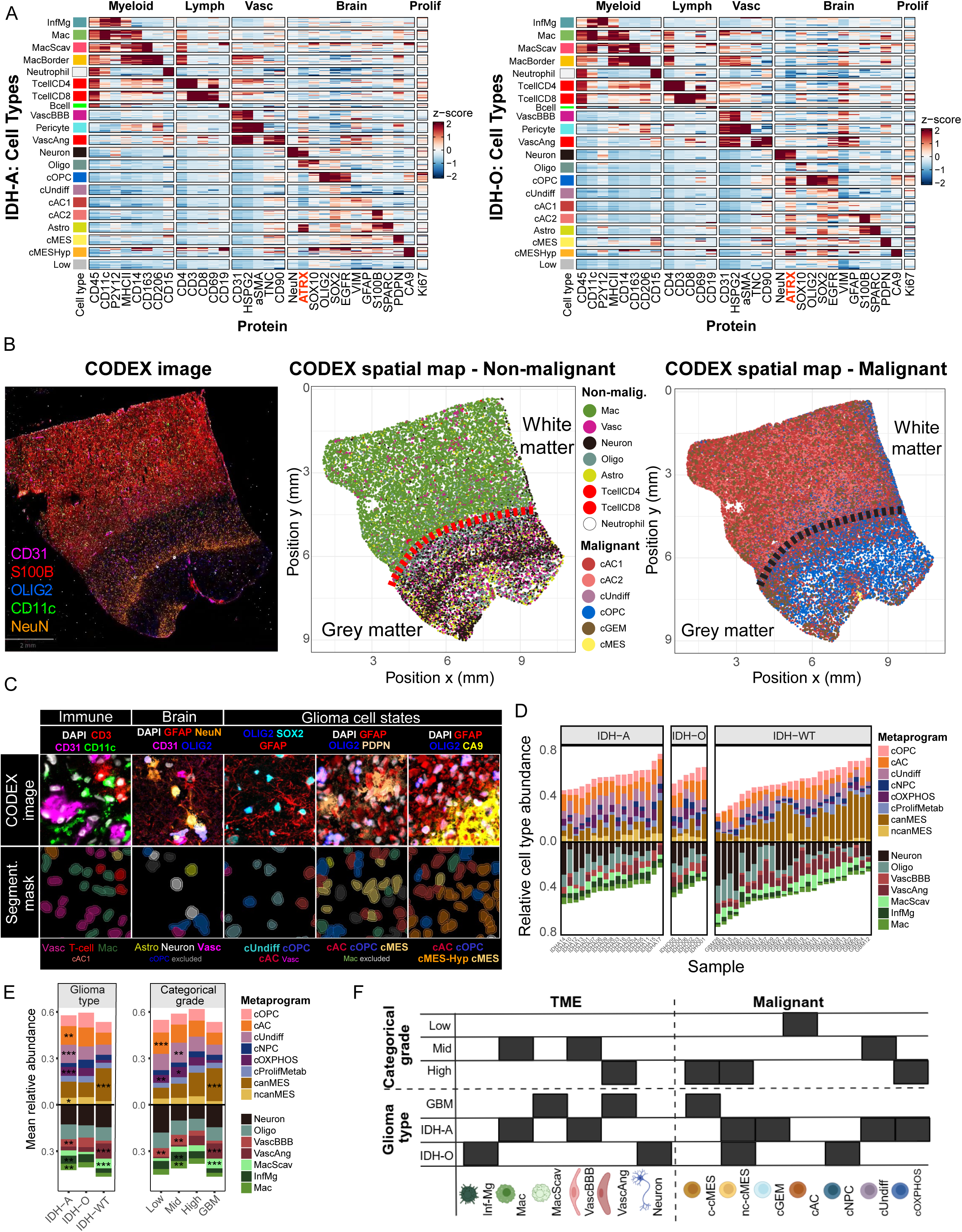
CODEX cell state annotation and compositional changes across glioma types and grades. **(A)** Heatmap of relative Nimbus scores (Z-score) per protein and cell type/state. Each row represents the indicated cell type or state in one sample, with varying numbers of rows across categories due to absence of certain populations in some samples. Heatmaps are stratified by glioma type (IDH-A vs. IDH-O) and highlight ATRX loss in malignant cellular states of IDH-A. **(B)** CODEX image of a sample showing the indicated markers (left), with corresponding spatial maps of annotated non-malignant TME cells (middle) and malignant cellular states (right). **(C)** Representative CODEX staining illustrating image segmentation. Top row, CODEX images with indicated markers; bottom row, corresponding to nuclear segmentation masks expanded by 2 µm and colored by cell state. Scale bar, 20 µm. **(D)** Bar plot showing relative abundance of Visium MPs per sample, stratified by glioma type. **(E)** Bar plot showing mean Visium MP abundance per glioma type and per categorial grade. **(F)** Heatmap showing significantly enriched cell types/states (black tiles) from CODEX or Visium data, grouped by categorial grade and glioma type.

Within the malignant compartment, clustering revealed the main glioma cell states cUndiff, cOPC, cAC, cMES, and cMES-Hyp. The antibody against IDH1 R132H was not compatible with the CODEX protocol and could therefore not be used to assign malignancy. However, consistent with pathology reports, all IDH-A samples showed the expected loss of ATRX in malignant cells, supporting the discrimination between malignant and non-malignant populations (**Fig. 2A**, **Fig. S2C**). In contrast, malignant cell states in IDH-O exhibited higher ATRX expression compared to their normal counterparts (e.g., cOPC vs. Oligo, cAC vs. Astro).

### Differences in tumor composition between glioma type and grade

All malignant MPs were detected across IDH-A, IDH-O, and GBM by Visium, but their relative abundances differed (**Fig. 2D-F**). IDH-A was enriched in cAC (*p =* 0.006), cOXPHOS (*p =* 0.0001), ncanMES (*p =* 0.04) and cUndiff (*p =* 0.0001), and GBM in canMES (*p =* 0.002). CODEX analysis further revealed differences in TME composition within *cellular tumor* regions based on IvyGAP classification (**Fig. 2F**, **Fig. S2D**): IDH-A contained fewer Neuron (0.5% vs. 2.5%, *p* = 0.001) and Vasc (4.2% vs. 6.7%, *p* = 0.03) but more Myeloid (14.7% vs. 4.7%, *p* = 0.005) than IDH-O. These patterns align with prior scRNA-seq and bulk RNA-seq studies^3^ and reflect the distinct anatomical niches of IDH-mut gliomas: IDH-O mostly residing in the neuronal- and vascular-rich cortical grey matter, and IDH-A in the myeloid-enriched subcortical white matter^2,14,15^.

Compositional analysis across categorical grades by Visium (**Fig. 2E,F**) revealed systematic changes in malignant and TME states across glioma grades. Low-grade gliomas were enriched in the cAC differentiated malignant states (*p* = 0.0003) and in cOXPHOS (*p* = 0.008). At intermediate grade, malignant populations shifted toward less differentiated programs (cUndiff, *p* = 0.002), accompanied by an increase in InfMg (*p* = 0.005), Mac (*p* = 0.003) and VascBBB (*p* = 0.008). At high-grade and GBM, tumors were dominated by hypoxia- and necrosis-associated malignant and non-malignant states (ncMES, VascAng, cMES, MacScav). Using CODEX, we further examined more granular TME changes. In IDH-A, myeloid cells shifted from inflammatory microglia (InfMg, P2Y12⁺) in low-grade to macrophages with an immunosuppressive phenotype (MacScav, CD163⁺, MacBorder, CD163⁺CD206⁺) in high-grade (*p* = 0.04; **Fig. S2E**). Similarly, the vascular compartment shifted from normal (VascBBB, CD31⁺GLUT1⁺) in low-grade IDH-A to aberrant vasculature (VascAng, CD31⁺GLUT1⁻TNC⁺) with increased ECM deposition in high-grade IDH-A (*p* = 0.04; **Fig. 2F**, **Fig. S2F**).

Taken together, these findings support a model in which low-grade IDH-mut gliomas are characterized by differentiated malignant states and a largely physiological microglia and vascular compartment. In intermediate grade, malignant cells become progressively de-differentiated, accompanied by expansion of Mac and VascBBB populations. Finally in high-grade IDH-mut gliomas, tumor composition shifts to hypoxia-associated malignant and non-malignant states that are more similar to GBM, including immunosuppressive macrophages and aberrant vasculature (VascAng).

### IDH-mut gliomas show low state-specific clustering compared to GBM

We next quantified glioma spatial organization using two complementary metrics: (i) *state-specific coherence* (State-Coh) and (ii) *consensus interactions*. State-Coh captures whether a specific state (defined by MP) forms coherent patches that are dominated by that state (State-Coh-high) or whether it is dispersed (State-Coh-low). We previously established this metric for Visium data as the fraction of neighboring spots dominated by the same MP (Methods, **Fig. S3A**)^9^.

IDH-A and IDH-O samples exhibited significantly lower State-Coh than GBM (*p* = 2.9×10⁻⁵, *p* = 2.1×10⁻⁶; **Fig. 3A**, **Fig. S3B**), indicating that IDH-mut gliomas generally adopt a State-Coh-low architecture. Across all glioma types, hypoxic samples showed significantly higher State-Coh than non-hypoxic samples (p = 3×10⁻⁵; **Fig. S3C**), consistent with hypoxia acting as a spatial organizing force when present in IDH-mut gliomas, as previously described in GBM^9^.

**Figure 3.**
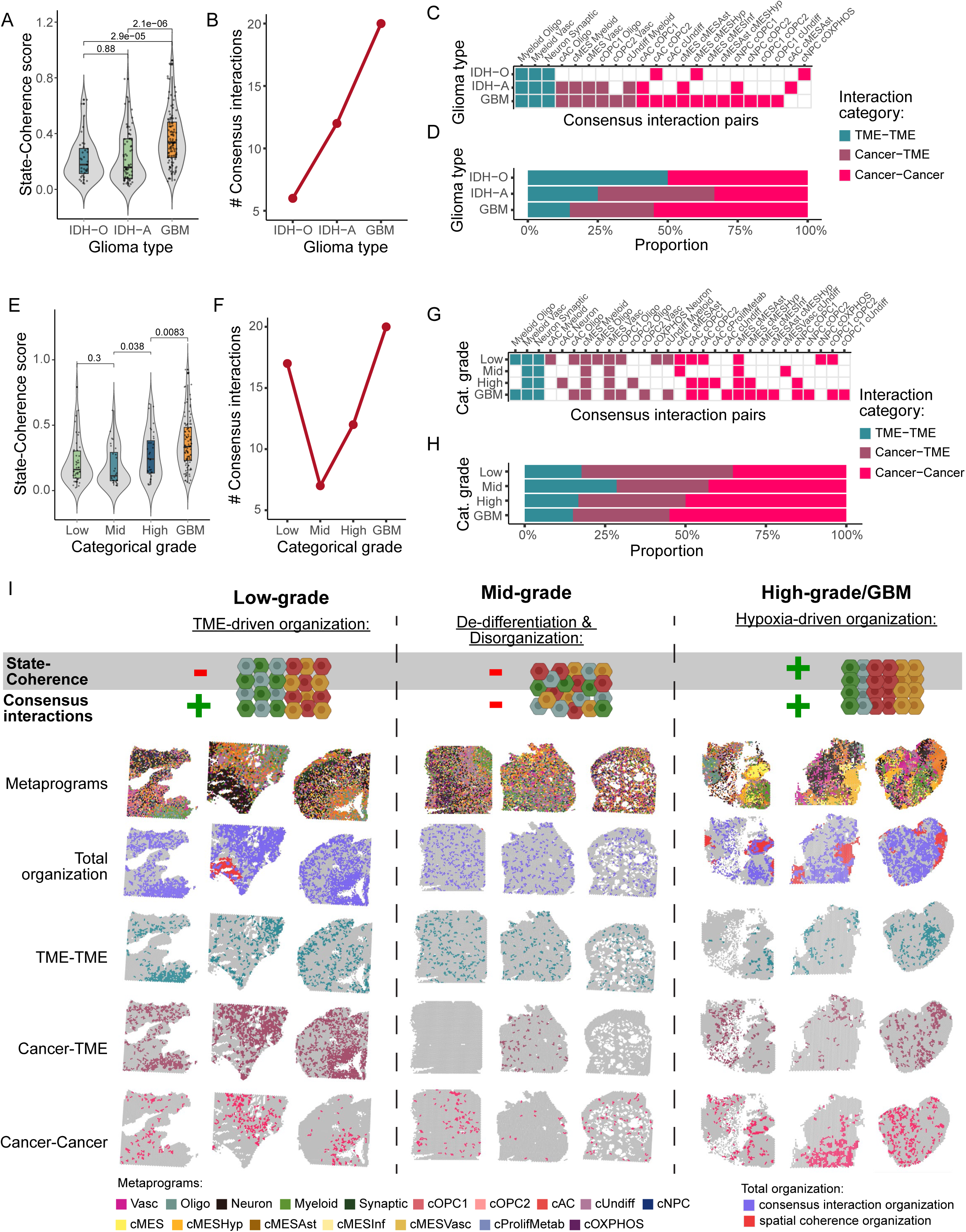
Three spatial archetypes of IDH-mut glioma. **(A)** Violin and boxplot of mean State-Coherence stratified by glioma type. *P* value*s* were computed using the Wilcoxon rank-sum test. **(B)** Line plot showing the number of consensus interactions per glioma type. **(C)** Heatmap of MP pairs engaged in consensus interactions for each glioma type, color-coded by interaction category (TME–TME, Cancer–TME, or Cancer–Cancer). **(D)** Proportion of interaction categories per glioma type. **(E)** Violin and boxplot of mean State-Coherence stratified by categorical grade. *P* values were computed using the Wilcoxon rank-sum test. **(F)** Line plots showing the number of consensus interactions (top) and consensus repulsions (bottom) per categorical grade. **(G)** Heatmap of MP pairs engaged in consensus interactions per categorical grade, color-coded by interaction category (TME–TME, Cancer–TME, or Cancer–Cancer). **(H)** Proportion of interaction categories per categorical grade. **(G)** Model of the three spatial archetypes, defined by the two spatial metrics: State-Coherence and consensus interactions. Representative Visium spatial maps are shown for three samples per category with five annotations: (*i*) MPs; (*ii*) Total organziation; (*iii*) TME–TME consensus interactions; (*iv*) Cancer–TME consensus interactions; and (*v*) Cancer–Cancer consensus interactions.

We next investigated spatial relationships between pairs of MPs by three methods spanning multiple scales of resolution (**Fig. S3D**, Methods)^9^. First, the degree of *colocalization* of two MPs was evaluated within the same Visium spot. Second, the *adjacency* of an MP pair was measured as the enrichment of one MP in spots within the immediate neighborhood of spots of the other MP. Third, we assessed the spatial association between MPs based on their *regional composition*, quantified as the correlation between their abundances within hexagonal windows of radius *r* (**Methods, Supplemental Note 1**). To focus on the most robust associations, we defined *consensus interactions* as those supported by more than one measure of spatial relationship and recurring across multiple samples (Methods). GBM was consensus interaction-rich (n = 20), followed by IDH-A (n = 12), whereas IDH-O was comparatively consensus interaction-poor (n = 6; **Fig. 3B**).

We next classified consensus interactions by partner identity (*interaction category*, Cancer–Cancer, Cancer–TME, and TME–TME; **Fig. 3C,D**). GBM was enriched in Cancer–Cancer couplings (*p* = 0.024; **Fig. S3G,H**), whereas IDH-O was enriched in TME–TME interactions (*p* = 0.003) and IDH-A was enriched in Cancer–TME interactions (*p* = 0.0013; Methods), highlighting the prominent role of microenvironmental coupling in IDH-mut gliomas.

### Low-grade IDH-mut gliomas have many consensus interactions

We next examined how these two spatial metrics vary across categorical grades. Low-grade IDH-mut gliomas were State-Coh-low (**Fig. 3E**), yet consensus interaction-rich (n = 17; **Fig. 3F**), with most couplings involving Cancer–TME and TME–TME pairs (*p* = 0.01; **Fig. 3G,H**, **Fig. S3I,J**). This pattern indicates recurrent cell-state pairings, possibly driven by brain anatomy even in the absence of the State-Coh-high organization typical of GBM.

In contrast, intermediate-grade gliomas were both State-Coh-low and consensus interaction-poor (n = 7; **Fig. 3F**), consistent with their highly diffuse nature and the disruption of normal brain structure. Two unique features of intermediate-grade gliomas were their significantly increased proportion of cUndiff within the malignant compartment (’Stemness score’, *p* = 0.009, **Fig. S3E**) and their increased malignant state diversity relative to other grades (*p* = 0.05, **Fig. S3F**), both of which may contribute to their lack of organization. In high-grade IDH-mut gliomas and in GBM, both State-Coh and consensus interactions increased significantly (State-Coh: *p* = 0.038, *p* = 0.008, respectively; consensus interactions: n = 12, n = 20; **Fig. 3E,F**), with a shift toward Cancer–Cancer couplings (NS), reflecting the emergence of hypoxia-associated global organization (**Fig. 3G,H**, **Fig. S3I,J**). The changes in number of consensus interactions across categorical grades were independently validated by high-resolution CODEX colocalization analyses (**Fig. S3K,L**) and were not explained by Visium cohort size differences (**Fig. S3M**).

To visualize the influence of the two organizational axes on overall tumor organization, we mapped onto representative Visium spatial maps the State-Coh-high regions and the spot-pairs that represent consensus interactions, which were further classified by interaction category (**Fig. 3I, Supplemental Note 2**). In low-grade tumors, Cancer–TME and TME–TME consensus interactions dominate despite overall State-Coh-low architecture. Intermediate-grade tumors are State-Coh-low and are consensus interaction-poor. In contrast, high-grade tumors and GBM are State-Coh-high and show a consensus interaction-rich architectures dominated by Cancer–Cancer interactions.

### The white-grey matter junction is a functional barrier for cell migration

In low-grade IDH-mut gliomas, the large fraction of TME-Cancer interactions may indicate a significant influence of physiological brain structures on tumor organization. While tumors establish a unique microenvironment that overrides many aspects of normal brain histology, it is still possible to identify remaining brain tissue compartments, such as the grey and white matter. We assigned 30% of the CODEX spatial data (from 19 tumors) into grey matter based on diffuse MAP2 and reduced PLP1 staining (Methods, **Fig. S4A**). Similarly, based on the opposite staining pattern, we assigned 70% of the CODEX data (from 23 tumors) to white matter.

We next focused on IDH-A (n = 7) and GBM (n = 6) samples, in which the cellular tumor is primarily located within the subcortical white matter (**Fig. 4A, S4B,C**). Four of the IDH-A tumors contained both white and grey matter regions, demarcated by sharp transitions of MAP2 and PLP1 staining, suggesting that the white–grey matter junction (WGJ) was maintained during glioma development (**Fig. 4A, S4A,B**). Notably, these junctions coincided with the transition from cellular to infiltrating tumor in all but one sample, suggesting that they serve as a functional barrier for cell migration. In 4 other IDH-A tumors, the transition from cellular to infiltrating tumor occurred within the white matter, offering a useful comparator spatial class of “white–white matter junctions” (WWJ, n = 4) (**Fig. S4C**). We therefore asked how glioma cells are distributed across these junctions and whether they are associated with changes in cell states.

**Figure 4.**
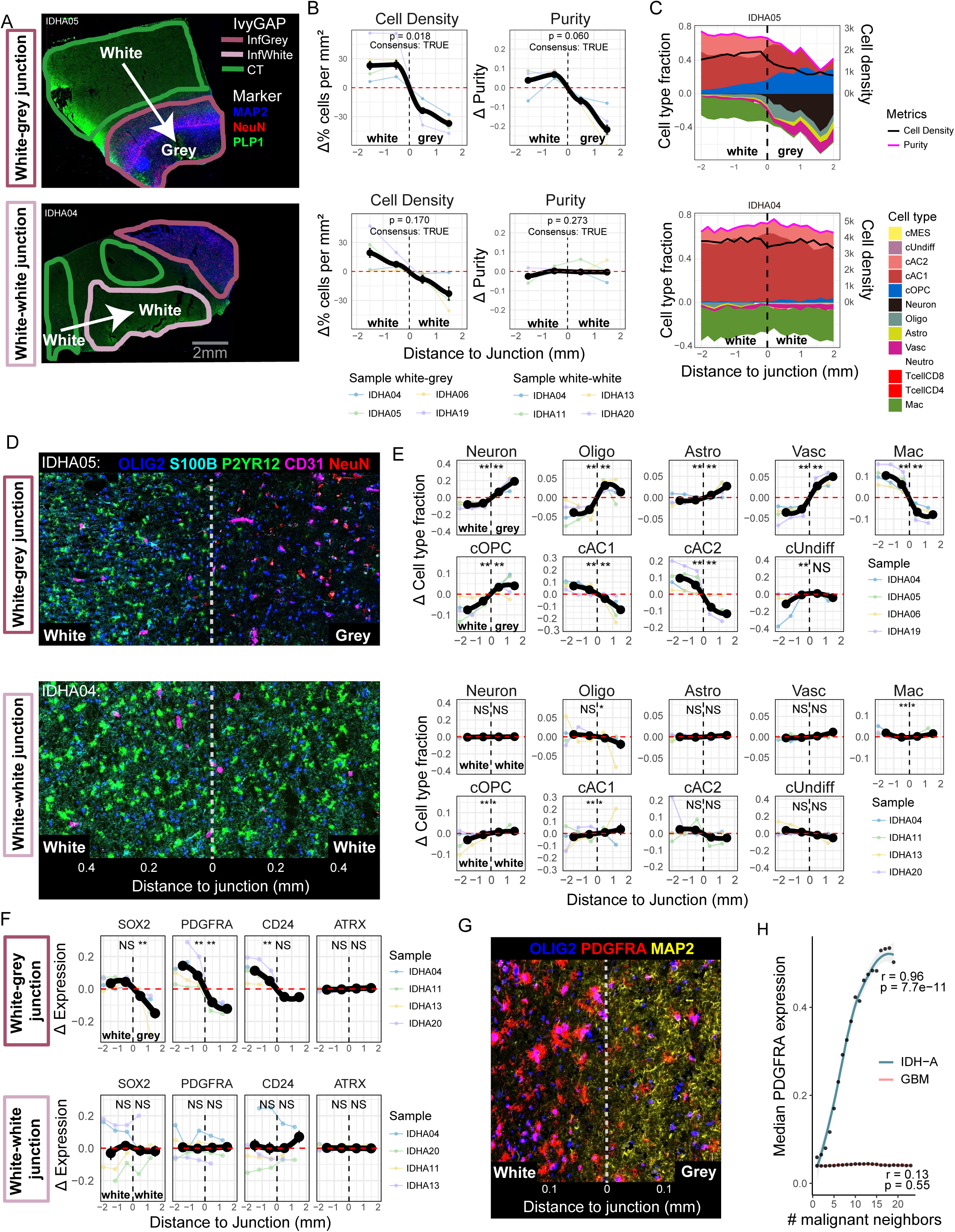
Barrier function of the white–grey matter junction. **(A)** Representative CODEX images of two samples (IDHA05, IDHA04) showing the indicated markers and overlaid by IvyGAP annotation. White arrows indicating the transition from the cellular tumor of white matter to infiltrating of grey matter (White-grey junction, top) and from the cellular tumor of white matter to infiltrating of white matter (White-white junction, bottom). **(B)** Line plot showing the relative changes of cell density and purity (malignant cell state fraction) at the white-grey junction (top) and the white-white junction (bottom). The black line represents the mean values of individual samples plotted in the background for 1mm bins. The red dashed line represents the mean of the two central bins. *P*-values are calculated using a paired *t*-test (Methods). Consensus was labeled TRUE if the directionality was identical in > 50% of the samples. **(C)** Area plots of the same samples (IDHA05, IDHA04) showing relative changes in cell-state fractions, cell density, and tumor purity across the white–grey junction (top) and white–white junction (bottom). Note the pronounced alterations at the white-grey junction compared with the white-white junction. **(D)** Representative CODEX images showing the indicated markers, highlighting cell-state composition differences at the white–grey junction (top) versus the white–white junction (bottom). **(E)** Line plots showing relative changes in cell-type fractions across the white–grey junction (top two rows) and the white–white junction (bottom two rows). Black lines represent mean values of individual samples (1-mm bins); the red dashed line indicates the mean of the two central bins. *P*-values were calculated using binomial tests on both sides of the junction, and directionality was required to be identical in > 50 % of samples (Methods). **(F)** Line plots showing relative changes in marker expression across the white–grey junction (top) and the white–white junction (bottom). Black lines represent mean values of individual samples (1-mm bins); the red dashed line indicates the mean of the two central bins. *P* values were calculated using two-sample Welch t-tests on both sides of the junction, with directionality consistent in > 50 % of samples (Methods). **(G)** Representative CODEX images showing the indicated markers, highlighting the decrease of PDGFRA expression in cOPC (OLIG2⁺) cell states after crossing into MAP2⁺ grey matter at the white–grey junction. **(H)** Scatter plot showing median PDGFRA protein expression as a function of the number of malignant neighbors within a 55-µm radius. PDGFRA expression in IDH-A samples is highly correlated with local malignant-cell density, whereas no correlation is observed in GBM.

We developed the *TmnApp* (Methods), which annotates cells based on their distance to a manually inserted reference line representing such junctions (**Fig. S4D**, **Movie S1**). We then analyzed the changes in cell density and composition as a function of their distance from the junctions. Interestingly, we observed two distinct behaviors depending on the transition type: a sharp decrease in cell density and tumor purity (i.e. malignant cell fraction) across the WGJ (**Fig. 4B, top panels**), in contrast to a more subtle and linear decrease across the WWJ (**Fig. 4B, bottom panels**). This suggests that WGJ acts as a functional barrier, preventing most glioma cells from crossing. The comparator WWJ may instead represent the margin of cellular tumor in this compartment, consistent with a gradual decrease in cell density and the absence of an associated microanatomic border.

### Changes in cell composition across the white-grey matter junction

Consistent with WGJ serving as a boundary, we observed not only a sharp global decrease in cell density, but also abrupt shifts in cell-state composition across the WGJ, which were consistent across the majority of cases (**Fig. 4C-E, Fig. S4E**). Non-malignant TME changes included the expected grey-matter enrichment of Neuron (0.8% vs. 28%, *p* < 0.001), Astro (0.1% vs. 4.4%, *p* < 0.001), and Vasc (3.5% vs. 10.7%, *p* < 0.001, **Fig. 4E**). In addition, there was an unexpected strong depletion of Mac in the grey-matter (23.1% vs. 1.2%, *p* < 0.001).

Among malignant states, cOPC cells were enriched after the WGJ (10.6% vs. 21.5%, *p* < 0.001), whereas cAC cells were excluded (AC1: 20.1% vs. 5.4%, *p* < 0.001, AC2: 15.2% vs. 0.8%, *p* < 0.001), suggesting that cOPC selectively crossed the barrier defined by the WGJ. No such effect was observed in GBM (**Fig. S5A,B**), consistent with the tendency of normal brain structures to erode in high grade gliomas (**Fig. S5C**).

The infiltrative behavior of cOPC mirrors the known migratory capacity of non-malignant OPCs in normal brain development and aligns with their identification as the key infiltrative state in GBM models in PDX mice^16^. We next examined whether cOPCs also exhibit phenotypic changes when crossing WGJ. Infiltrating cOPCs in the grey-matter showed increased expression of canonical Oligo marker (SOX10) and the astrocytic marker SOX9. The expression of the stemness marker SOX2 was decreased and PDGFRA was lost, without changes in ATRX (**Fig. 4F, Fig S5D**). These changes are consistent with increased differentiation of malignant cOPCs into an Oligo-like phenotype after crossing. Interestingly, the PDGFRA loss of cOPC in the grey matter, correlated remarkably well with malignant cell density in IDH-mut gliomas (*r =* 0.96, *p* = 7.7e-11) but not in GBM (*r =* 0.13, *p* = 0.55, (**Fig. 4G,H**).

In summary, we identified an unexpected barrier function of the WGJ, in which the few cells that do cross this barrier are strongly enriched for cOPC, and they tend to exhibit a more differentiated Oligo-like state (**Fig. 5**).

**Figure 5.**
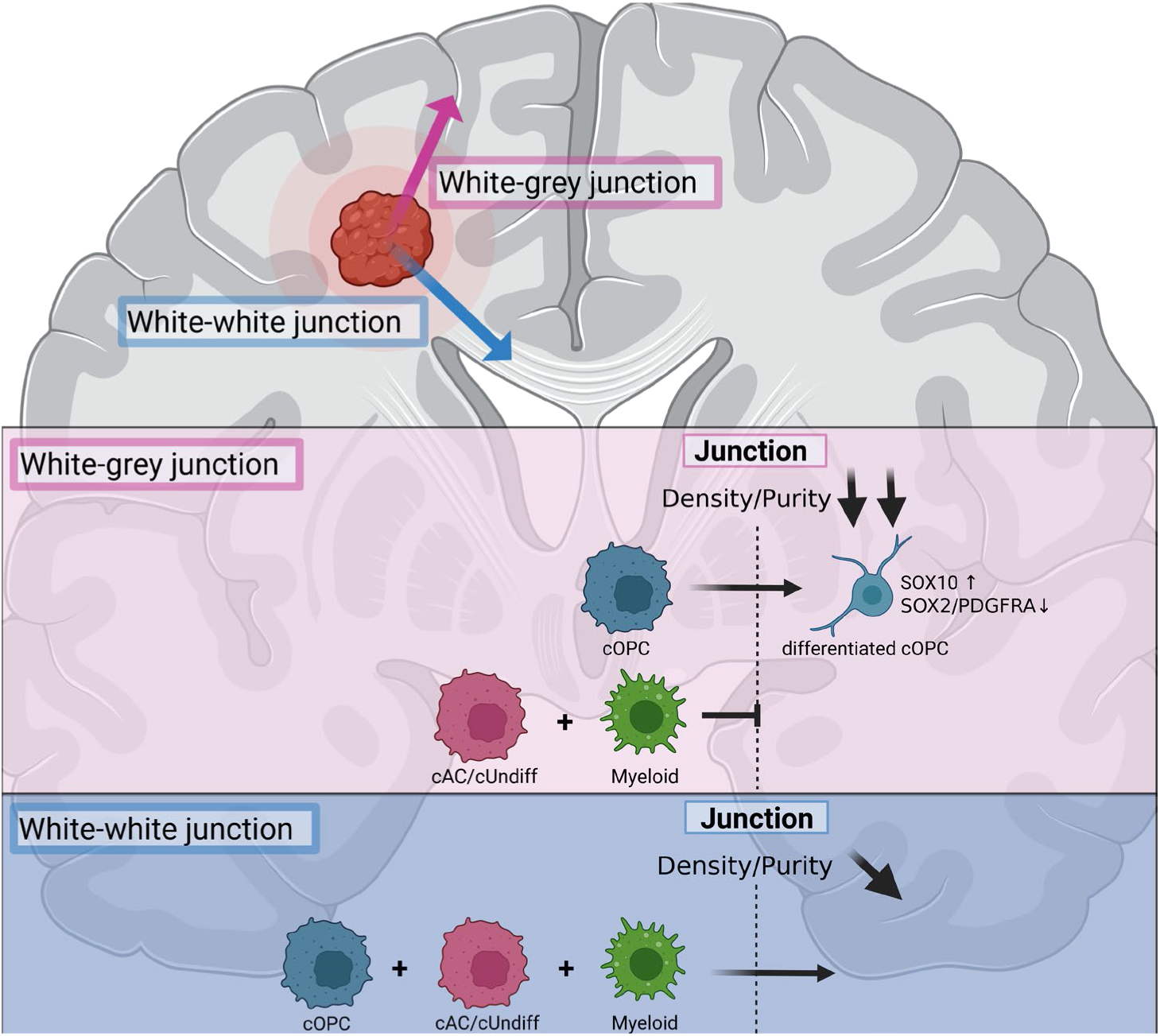
Compartment-dependent model of glioma invasion. Schematic summary illustrating differences in invasive behavior across the white–grey junction compared with the white–white junction.

## Discussion

### Quantifying tumor organization

We combined spatial transcriptomics and spatial proteomics to map the organization of IDH-mut gliomas across histological grades using two complementary metrics: (i) *State-Coh*, which quantifies how clustered versus scattered a specific cell type or state is within the tissue, and (ii) *consensus interactions*, defined as cell–cell couplings that occur significantly more often than expected by chance, persist across multiple spatial scales, and recur across samples. State-Coh-high implies the appearance of small tumor regions (i.e. patches) that are each dominated by a single cell type or cell state, while a high number of consensus interactions (consensus interaction-rich) implies that specific pairs of cell types/states tend to be proximal and thereby have a larger impact on tumor organization.

In our previous work on GBM, State-Coh-high regions surrounding hypoxia and necrosis were also consensus interaction-rich, suggesting that the two metrics may be related^9^. Here, we find that low-grade IDH-mut gliomas break this association: they are consensus interaction-rich despite being State-Coh-low, indicating that these metrics capture complementary dimensions of tissue organization. Across the IDH-mut cohort, these two axes reveal three organizational archetypes aligned with grade (**Fig. 3I, Supplemental Note 2**): (i) low-grade—consensus interaction-rich but State-Coh-low; (ii) intermediate-grade— consensus interaction-poor with only the presence of universal consensus interactions (i.e. Myeloid-cMES, Vasc-Myeloid) and State-Coh-low; and (iii) high-grade—consensus interaction-rich and State-Coh-high. The latter archetype mirrors the layered, hypoxia-anchored architecture previously described in GBM, where necrosis and hypoxia impose long-range constraints that organize malignant and microenvironmental states into defined spatial layers. In contrast, the early archetype at least partially reflects the impact of residual brain parenchymal structures, as discussed further below. Finally, the intermediate archetype appears to reflect the absence of strong influences from either hypoxia/necrosis or parenchymal brain structures, in which case cellular plasticity and perhaps cellular mobility may give rise to largely disorganized tumors.

### A mechanistic model for the three spatial archetypes

We propose three spatial archetypes that capture how glioma organization changes across grades. Because IDH-mut gliomas typically progress in a stepwise manner, categorical grades can serve as a surrogate for this trajectory and thus offer an opportunity to infer organizational changes during progression without longitudinal sampling.

To interpret these archetypes, it is essential to consider the structured nature of the environment in which these tumors arise. The human brain is intrinsically structured: grey matter is made of dense neuronal circuits supported by astrocytes and other glia, while white matter features long axonal tracts myelinated by oligodendrocytes^17^. During glioma genesis, malignant cells arise within this highly organized microenvironment and must interact with it. Consistent with this, low-grade IDH-mut gliomas exhibit short-range consensus couplings between malignant and non-malignant cell states (e.g., cOPC–Oligo, cOXPHOS–Neuron, cMES–Myeloid; **Fig. 3G**). We have reported such interactions previously: myeloid cells can induce MES-like states via the OSM–OSMR axis^18^, and oligodendrocytes are spatially coupled with cOPC-like programs in GBM^9^. These interactions manifest at micron scales without clustering of individual cell types (State-Coh-low), resulting in a disorganized “salt-and-pepper” pattern despite recurring local coupling.

As tumors advance to intermediate-grade, malignant cells increase their proliferation and de-differentiation (**Fig. 2F, Fig. S3E**)^3,6^ accompanied by an almost complete collapse of consensus interactions (**Fig. 3F**). We hypothesize that this phase reflects increased malignant plasticity together with progressive destruction of physiological brain architecture. The latter is supported by a recent study demonstrating that radiological and histological disruption of white-matter tracts correlates with grade in IDH-mut gliomas^19^, and by *in-vivo* models showing that glioma cells can induce axonal injury and Wallerian-like degeneration, thereby amplifying neuroinflammation and promoting progression^20^.

In high-grade gliomas, including GBM, hypoxia and necrosis emerge as dominant constraints that impose organization: State-Coh increases across malignant and TME compartments (**Fig. 3E, Fig. S3C**), and Cancer–Cancer consensus interactions prevail (**Fig. S3I,J**). Distinct cell state clusters organize into layers anchored by pseudopalisading necrosis, recapitulating the canonical hypoxia-driven microenvironment that also contains microvascular proliferation^2,9,10^.

Interestingly, these observed spatial archetypes align well with theoretical models proposed previously for multicellular tissue organization^21,22^. These models predicted that, in the absence of extrinsic gradients, tissues would adopt salt-and-pepper configurations (as we observe in early progression), while the introduction of external forces, such as hypoxia, would yield islet- or stripe-like domains (as we observe in late progression and GBM).

### The barrier function of the white–grey matter junction

Building on this framework, we next examined how physiological brain structures can influence glioma organization. Here, the white-grey matter junction (WGJ) emerged as a striking anatomical landmark that shapes the spatial distribution of both malignant and non-malignant compartments in early- and mid-progression gliomas. We found that the characteristic myeloid inflammation of gliomas^10,18,23^ is largely confined to the white matter, with very few myeloid cells in the grey matter (**Fig. 4E, Fig. S4E**). Likewise, most malignant cells were excluded from the cortical side of the junction—with the notable exception of cOPC, which may preferentially traverse this interface.

Given the diffuse growth of IDH-mut gliomas, we were surprised to find that the WGJ appears to be a significant anatomic boundary for malignant cell infiltration in low-grade tumors. The WGJ may serve as a physical barrier to glioma infiltration that restricts infiltration into the cortex. Alternatively, the white matter itself may serve as a more permissive environment for malignant cell growth and migration, beyond which adaptation – and associated fitness costs – are required for tumor cells that successfully infiltrate the cortex. Our data suggests that cOPCs exhibit a selective advantage relative to other tumor cell states in crossing this boundary. cOPC displayed phenotypic changes consistent with such adaptation, including downregulation of the stemness marker SOX2 and other changes that may suggest the acquisition of a more specific differentiated state. These findings align with prior work showing that cOPC and vasculature are spatially coupled^9^ and might reflect the hijacking of a developmental OPC-like migratory program that exploits perivascular rails^24^ to infiltrate surrounding brain tissue.

The ability of cOPC to migrate along blood vessels may enable them to circumvent non-cellular constraints defining the junction. The grey-matter ECM is denser and enriched in sulfated proteoglycans, including chondroitin sulfate proteoglycans (CSPGs) such as neuroglycan C (CSPG5) (**Fig. S5E,F**)^25^. In addition, the tissue architecture transitions sharply from the anisotropic, aligned white-matter tracts to a more isotropic, randomly oriented cortical ECM, altering both topography and mechanical properties^26^. Both the compositional and mechanical discontinuities likely contribute to the barrier-like properties of the WGJ, whereas perivascular migration offers a direct path that traverses this boundary^27^.

In summary, we provide an extensive spatial description of IDH-mut glioma and define organizational architypes of tumor progression. The quantitative framework established here can be applied to other cancers to test whether progression follows reproducible spatial design principles, and whether therapeutic perturbations rewire these organizational rules.

### Limitations of the study

This study has several limitations. First, given the large number of tumor categories (glioma type, grade) the number of tumors per category remains modest. We focused on obtaining large, intact tissue sections—as opposed to tissue microarrays (TMAs) with more samples—to preserve the larger spatial context necessary to capture higher-order tumor organization across progression stages. Second, the spatial resolution of spot-based Visium limits single-cell discrimination, which we mitigated through orthogonal validation with high-resolution CODEX imaging.

## Acknowledgements

This work was supported by the European Research Council Consolidator Grant and an Israel Science Foundation grant (to I.T.). I.T. is the incumbent of the Dr. Celia Zwillenberg-Fridman and Dr. Lutz Zwillenberg Career Development Chair, and is supported by the Zuckerman STEM Leadership Program. The study was also supported by The Mark Foundation Emerging Leader Award (M.L.S.), the V Foundation All-Star Translational Award (M.L.S.), N.I.H. R37CA245523 and R01CA258763 (both to M.L.S.). This study was also supported by the Deutsche Forschungsgemeinschaft (DFG, German Research Foundation) Project-ID 441891347—SFB1479 (R.H.) and by the Mertelsmann Foundation (R.H.). R.H. is funded by the IMMediate Advanced Clinician Scientist-Program, Department of Medicine II, Medical Center – University of Freiburg and Faculty of Medicine, University of Freiburg, funded by the Federal Ministry of Research, Technology and Space (Bundesministerium für Forschung, Technologie und Raumfahrt, BMFTR) - 01EO2103. N.G.D. is supported by the Israeli Council for Higher Education (CHE) via the Weizmann Data Science Research Center and by a research grant from the Estate of Tully and Michele Plesser. V.M.R. is supported by a NIH T32 Cancer Neuroscience Training Grant (T32CA272386). V.M. is supported by a NIH T32 Cancer Neuroscience Training Grant (T32CA272386). This work was supported by BMBF (DiaQNOS: 1010 0665 01) (J.B.)and DFG MA 10605/1- 666 1 (V.M.R., K.J.). The images in this paper were acquired at the Advanced Optical Imaging Unit, de Picciotto-Lesser Cell Observatory Unit at the Moross Integrated Cancer Center Life Science Core Facilities, Weizmann Institute of Science. The authors would like to thank Abcam for the generous donation of carrier-free antibodies for CODEX experiments.

## Author Contributions

R.H., A.C.G., N.G.D., C.W.M. and I.T. conceived the project, designed the study, interpreted results, and wrote the manuscript. M.C.N., and M.L.S. provided guidance, feedback, and edited the manuscript. R.H., A.C.G., and N.G.D. performed computational analyses and together with D.S. developed computational methods for spatial analysis. L.N.G.C., M.C.N., A.B.P, J.B. and M.L.S. provided glioma samples for spatial transcriptomics and spatial proteomics. V.M.R. and K.J. provided spatial transcriptomics data of additional samples. R.H. performed spatial proteomics and tissue sectioning. Y.T. generated the brain compartment classifier. A.C.G. and C.W.M. performed spatial transcriptomics. C.W.M. annotated histology sections. I.G., O.G. and Y.A. provided imaging and microscopy expertise and CODEX support. M.K. and H.K.S. provided genomics expertise and Visium support. I.T. and M.L.S. supervised the study.

---

**Figure S1.**
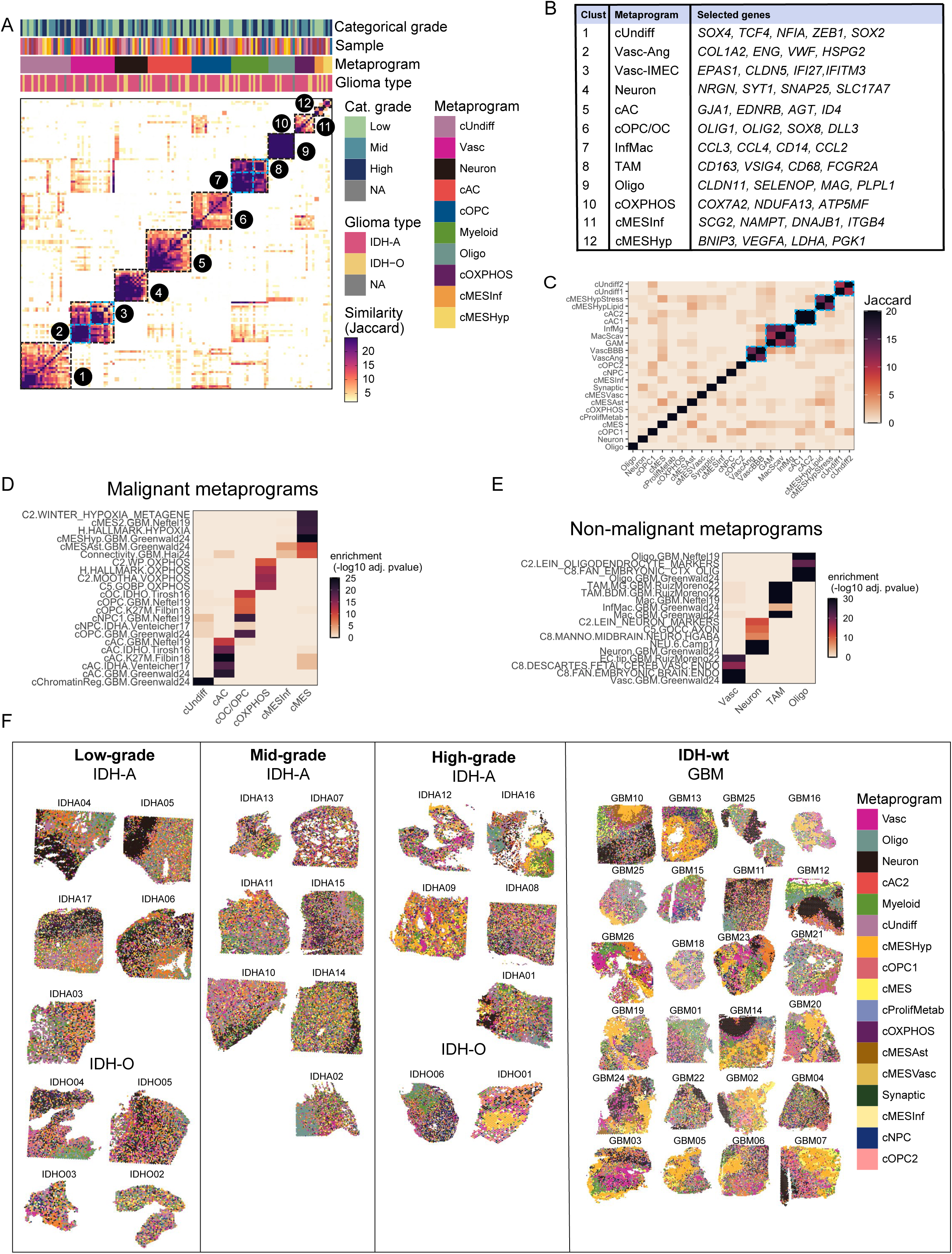
Metaprogram generation and Visium spatial maps, related to Figure 1. **(A)** Similarity matrix based on gene identity overlap quantified by Jaccard index, for all programs derived from NMF and Leiden clusters from IDH-mutant only samples. Cluster numbers correspond to those in the table in (B). **(B)** Table of spatial IDH-mutant metaprogram names and selected genes corresponding to clusters numbered in (A). **(C)** Jaccard heatmap of all glioma metaprograms. Highly overlapping gene expression programs of the same cell state or same cell type (highlighted in blue) are collapsed to a single program for downstream spatial analysis. **(D)** Enrichments of malignant spatial metaprograms (rows) with gene-sets that were previously defined (columns). Enrichment calculated by hypergeometric test (-log10 FDR-adjusted p-values). **(E)** Enrichments of non-malignant spatial metaprograms (rows) with gene-sets that were previously defined (columns). Enrichment calculated by hypergeometric test (-log10 FDR-adjusted p-values). **(F)** Spatial maps of Visium samples with spots annotated by dominant metaprogram. Spots were scored for metaprograms and annotated by their maximum score.

**Figure S2.**
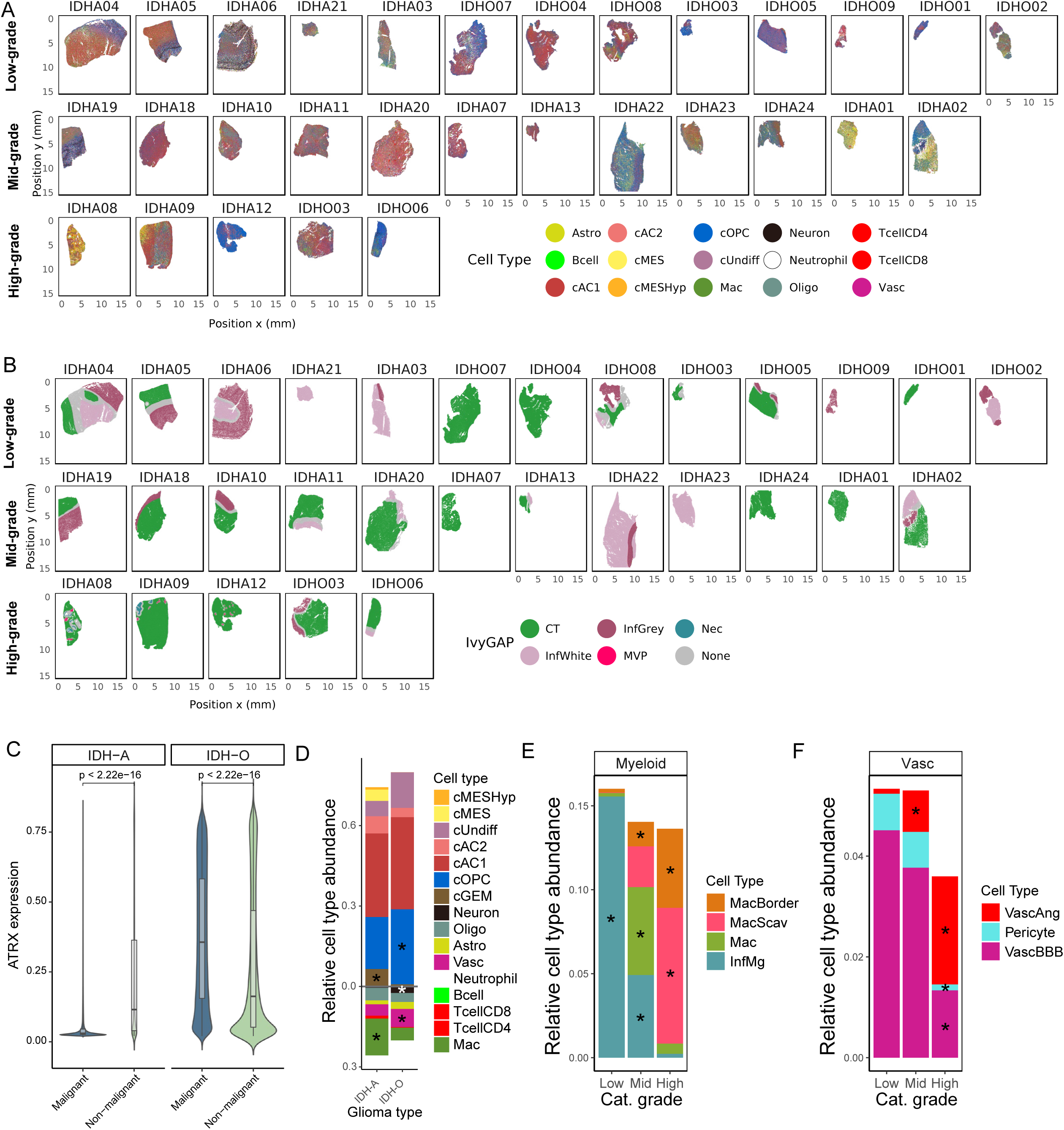
CODEX Spatial maps and sample compositions, related to Figure 2. **(A)** Spatial maps of CODEX samples with single cells annotated by cell type or state. **(B)** Spatial maps of CODEX samples with regional annotations by IvyGAP. **(C)** Violin plot showing the ATRX expression of malignant vs. non-malignant cells in IDH-A and IDH-O. **(D)** Bar plot showing relative cell type abundance per glioma type. Significance by Wilcoxon-rank sum test, p-value<0.05 *. **(E)** Bar plot showing relative myeloid cell type abundance per categorical grade. Significance by Wilcoxon-rank sum test, p-value<0.05 *. **(F)** Bar plot showing relative vasculature cell type abundance per categorical grade. Significance by Wilcoxon-rank sum test, p-value<0.05 *.

**Figure S3.**
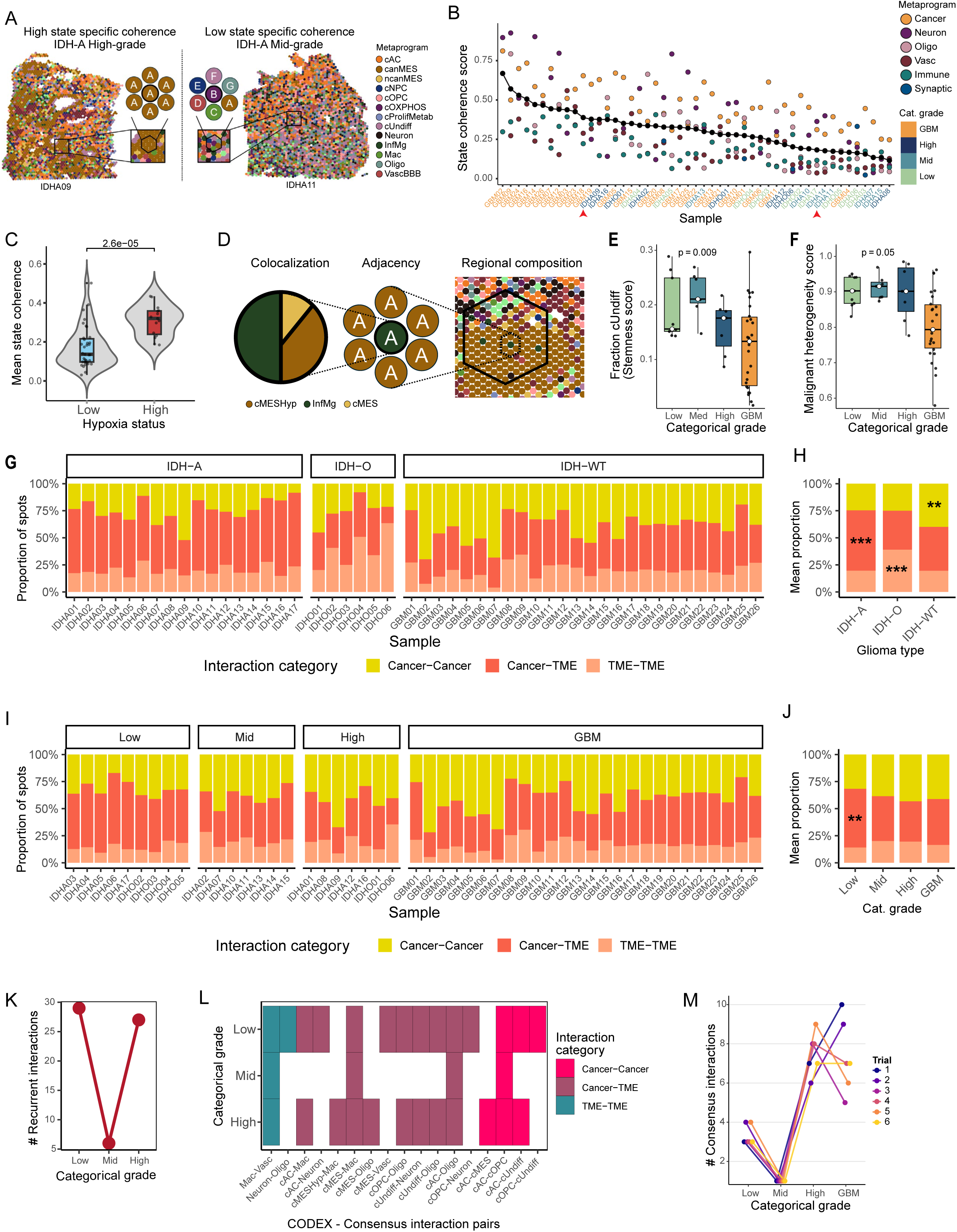
State coherence and consensus interactions, related to Figure 3. **(A)** Spatial map annotated by metaprograms for IDH09 (high grade) and IDH11 (intermediate-grade) illustrating high state-specific coherence and low state-specific coherence. **(B)** State-specific coherence by sample. Malignant MPs are grouped together for calculation. Red arrows point to the samples shown in Figure S3A. **(C)** Violin plot of mean state coherence across all MPs, for samples with high abundance (>10%) of cMESHyp vs. low abundance of cMESHyp. *P*-value is shown on the plot (by Wilcoxon rank-sum test). **(D)** Scheme illustrating spatial association measures between pairs of metaprograms across scales from high to low resolution. **(E)** Box plot of per sample “Stemness” score by categorical grade (i.e., the proportion of “cUndiff” within the malignant compartment of each sample). Significance was calculated by Wilcoxon rank-sum test; *p*-value = 0.0092 for intermediate-grade. **(F)** Box plot of per sample “Malignant heterogeneity” score by categorical grade (i.e., normalized Shannon entropy of malignant MPs). Significance was calculated by Wilcoxon rank-sum test; *p*-value = 0.0488 for intermediate-grade. **(G)** Interaction category composition (Cancer-Cancer, Cancer-TME, TME-TME) among “allowed pairs” per sample between tumor types (see Methods for definition of “allowed pairs”). Significance by Wilcoxon-rank sum test, FDR-adjusted p-value < 0.05 *, 0.01**, 0.001***. **(H)** Summary of interaction category composition shown in (G) (mean proportions for each type), statistics as defined in (G). **(I)** Interaction category composition (Cancer-Cancer, Cancer-TME, TME-TME) among “allowed pairs” per sample between categorical grades (see Methods for definition of “allowed pairs”). Significance by Wilcoxon-rank sum test, FDR-adjusted p-value<0.05 *, 0.01**, 0.001***. **(J)** Summary of interaction category composition shown in (I) (mean proportions for each type), statistics as defined in (I). **(K)** Line plot showing the number of recurrent interactions by categorical grade assessed with the colocalization metric (Methods). **(L)** Heatmap of CODEX Cell type/state pairs engaged in recurrent interactions per categorical grade as defined in (K). Color-code by interaction category (TME–TME, Cancer–TME, or Cancer–Cancer). **(M)** Resampling control for robustness of recurrent interactions. Each categorical grade was down sampled to a common sample size (n=6) for six independent trials. For each trial, adjacency analysis was performed and the number of consistent enriched interactions across samples within each categorical grade was computed.

**Figure S4.**
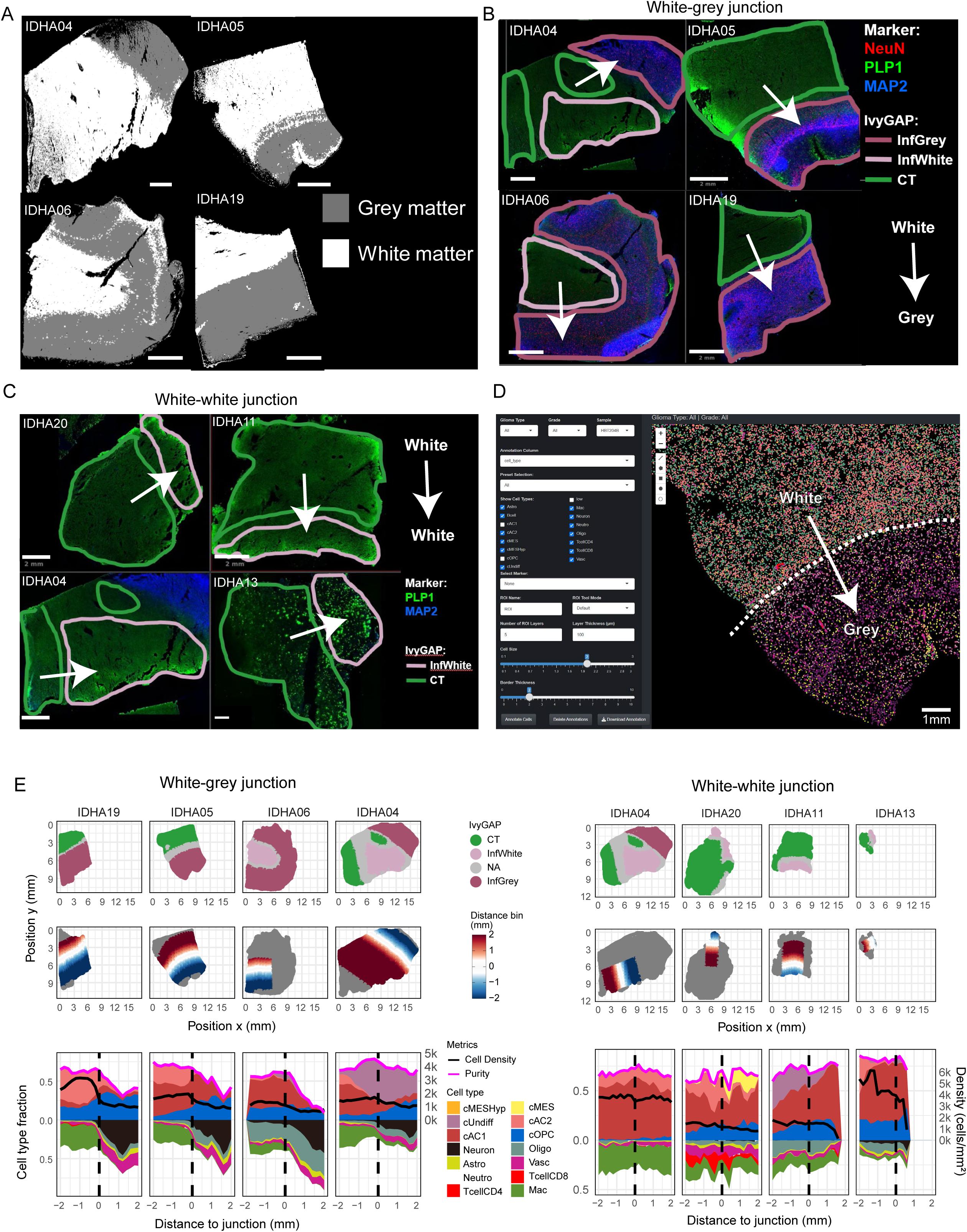
CODEX junction analysis of IDH-mut gliomas, related to Figure 4. **(A)** Spatial maps showing CODEX samples after segmentation of the white- and grey matter using PLP1 and MAP2 expression (Methods). Scale bars, 2mm. **(B)** CODEX images of samples containing white-grey junction with overlay of IvyGAP annotation. The indicated markers are shown. The white arrow crosses the junction from white- to grey-matter. Scale bars, 2mm. **(C)** CODEX images of samples containing white-white junction with overlay of IvyGAP annotation. The indicated markers are shown. The white arrow crosses the junction from white matter with cellular tumor to infiltrated white matter. Scale bars, 2mm. **(D)** Screenshot of the spatial annotation application “TmnApp”, showing the interactive functionality. The dashed white line shows the white-grey junction, that serves as reference line for distance annotation of all cells. **(E)** Spatial maps of CODEX samples with white-grey (left) or white-white junction (right) annotated by IvyGAP (top row) or distance to junction (middle row). The bottom row shows area plots of the respective samples visualizing the cell type fraction changes as function of distance to the junction. The black line shows cell density and the pink line purity (i.e. malignant fraction of cell types/states.

**Figure S5.**
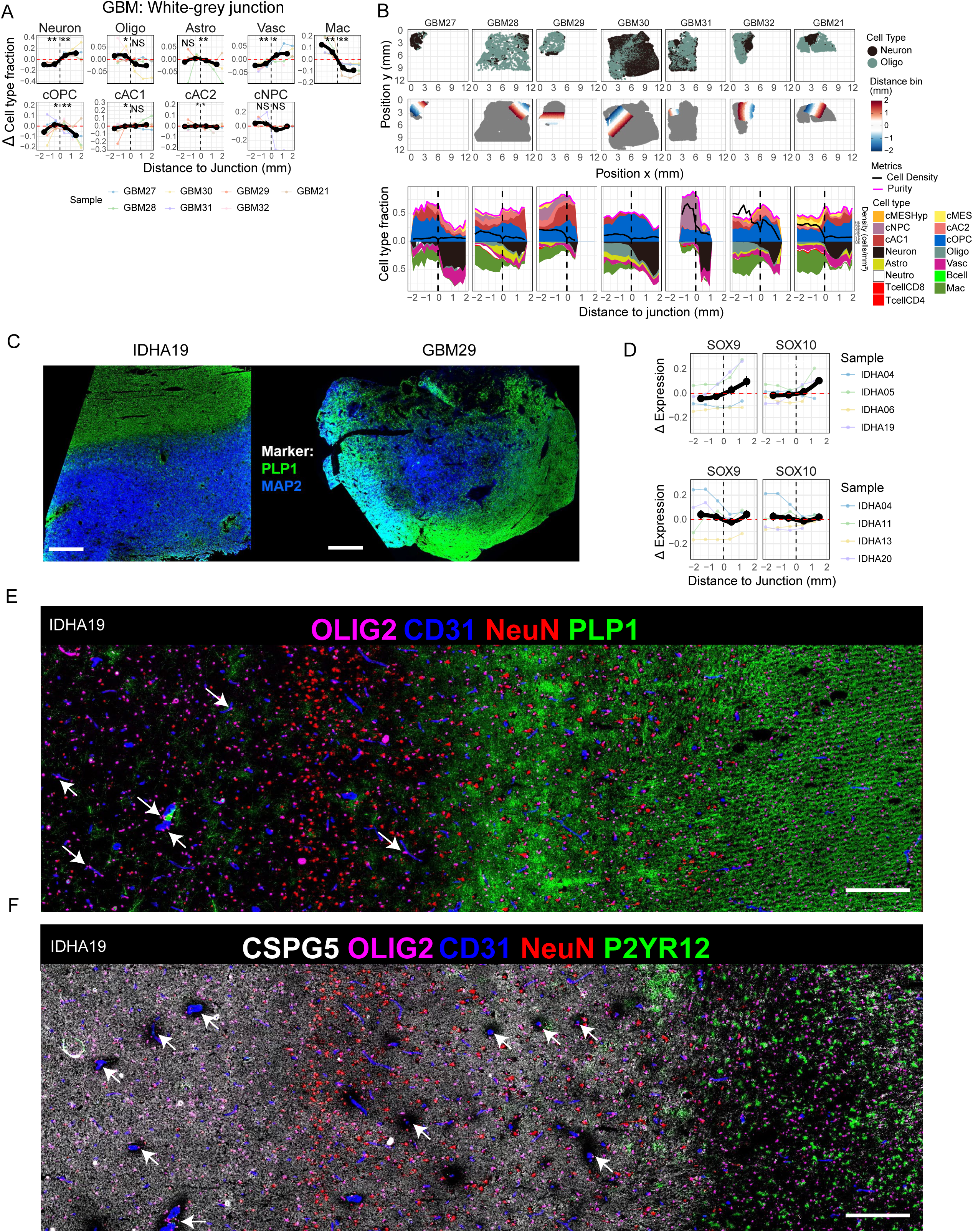
Characteristics of the white-grey junction, related to Figure 5. **(A)** Line plots showing relative changes in cell-type fractions across the white–grey junction in GBM. Black lines represent mean values of individual samples (1-mm bins); the red dashed line indicates the mean of the two central bins. *P*-values were calculated using binomial tests on both sides of the junction, and directionality was required to be identical in > 50 % of samples (Methods). **(B)** CODEX spatial maps of GBM samples with white-grey junction annotated by Neuron and Oligo (top row) or distance to junction (middle row). The bottom row shows area plots of the respective samples visualizing the cell type fraction changes as function of distance to the junction. The black line shows cell density and the pink line purity (i.e. malignant fraction of cell types/states. **(C)** CODEX image for the indicated markers showing a representative IDH-A sample (left) with a sharp white-grey junction. In contrast, the white-grey junction in GBM (right) looks diffuse and eroded. Scale bars, 2mm. **(D)** Line plots showing relative changes in marker expression across the white–grey junction (top) and the white–white junction (bottom) in IDH-A. Black lines represent mean values of individual samples (1-mm bins); the red dashed line indicates the mean of the two central bins. *P*-values were calculated using two-sample Welch *t*-tests on both sides of the junction, with directionality consistent in > 50 % of samples (Methods). **(E)** CODEX image of the white–grey matter junction in sample IDHA19 showing the indicated markers. Numerous cOPC cells (OLIG2⁺) are in close contact with vasculature (CD31⁺), suggesting migration along vascular structures (white arrows). Scale bar, 200 µm. **(F)** CODEX image of the same region shown in **(E)** displaying the indicated markers. White arrows highlight vascular structures containing cOPCs within the grey matter that are spared from dense extracellular matrix components, including neuroglycan-C (CSPG5). Scale bar, 200 µm.

**Supplemental Note 1.**
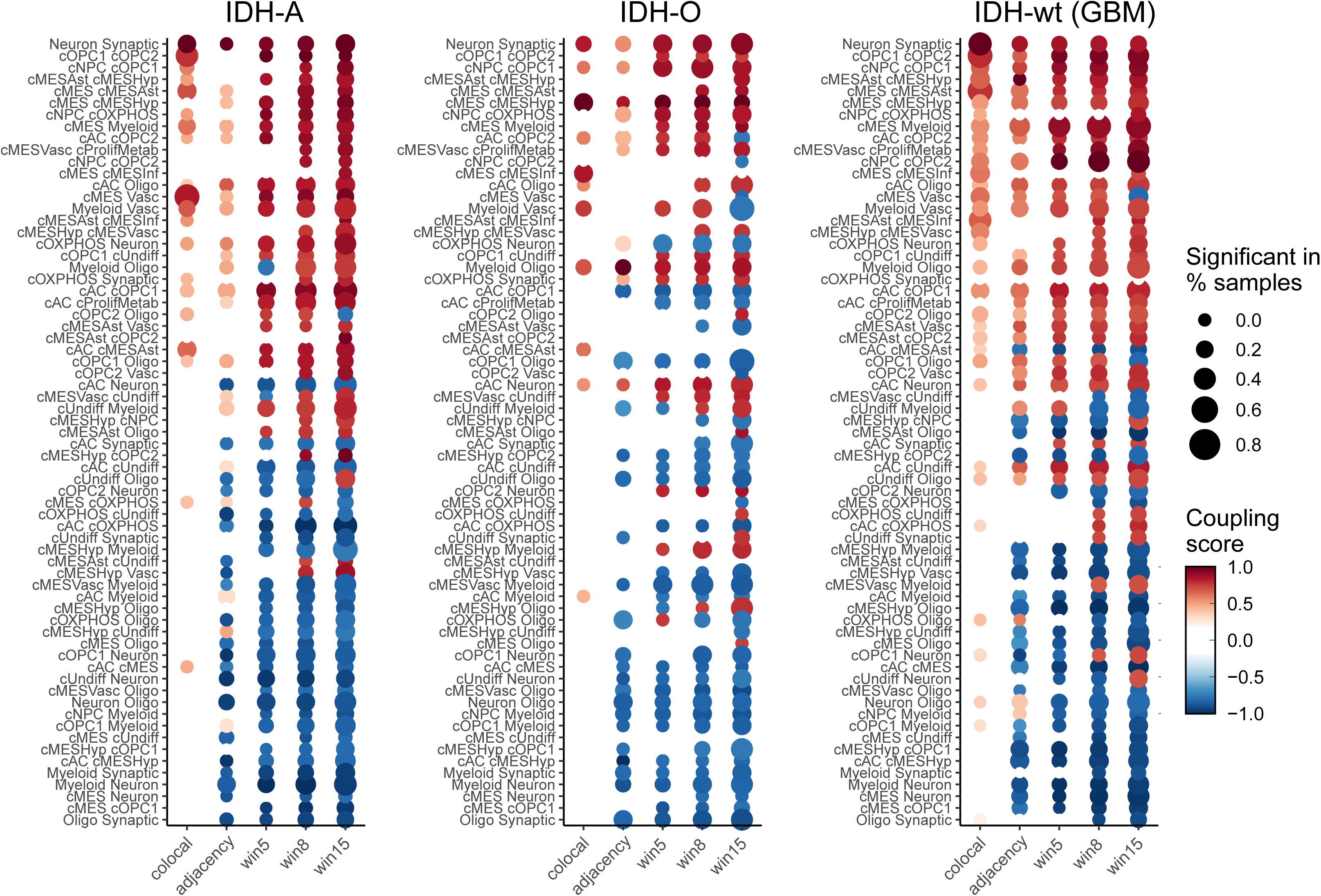
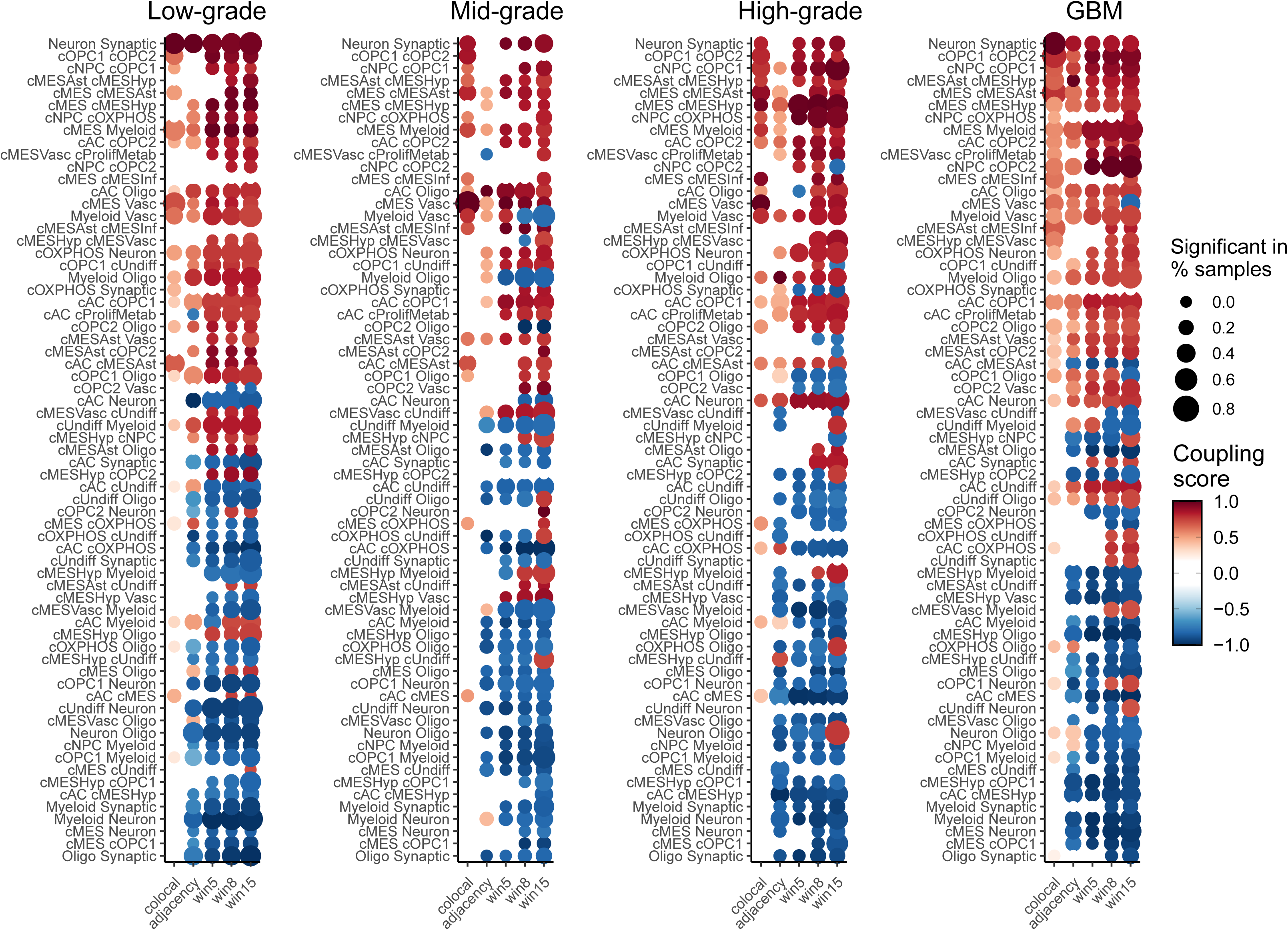
Heatmap of MPs spatial interaction across scales. Columns show the scores of different spatial relationship measures across all samples from the same type or categorical grade. Dots are colored by mean scaled interaction strength, and dot size corresponds to the fraction of samples in which the relationship is significant by Fisher’s exact test.

**Supplemental Note 2.**
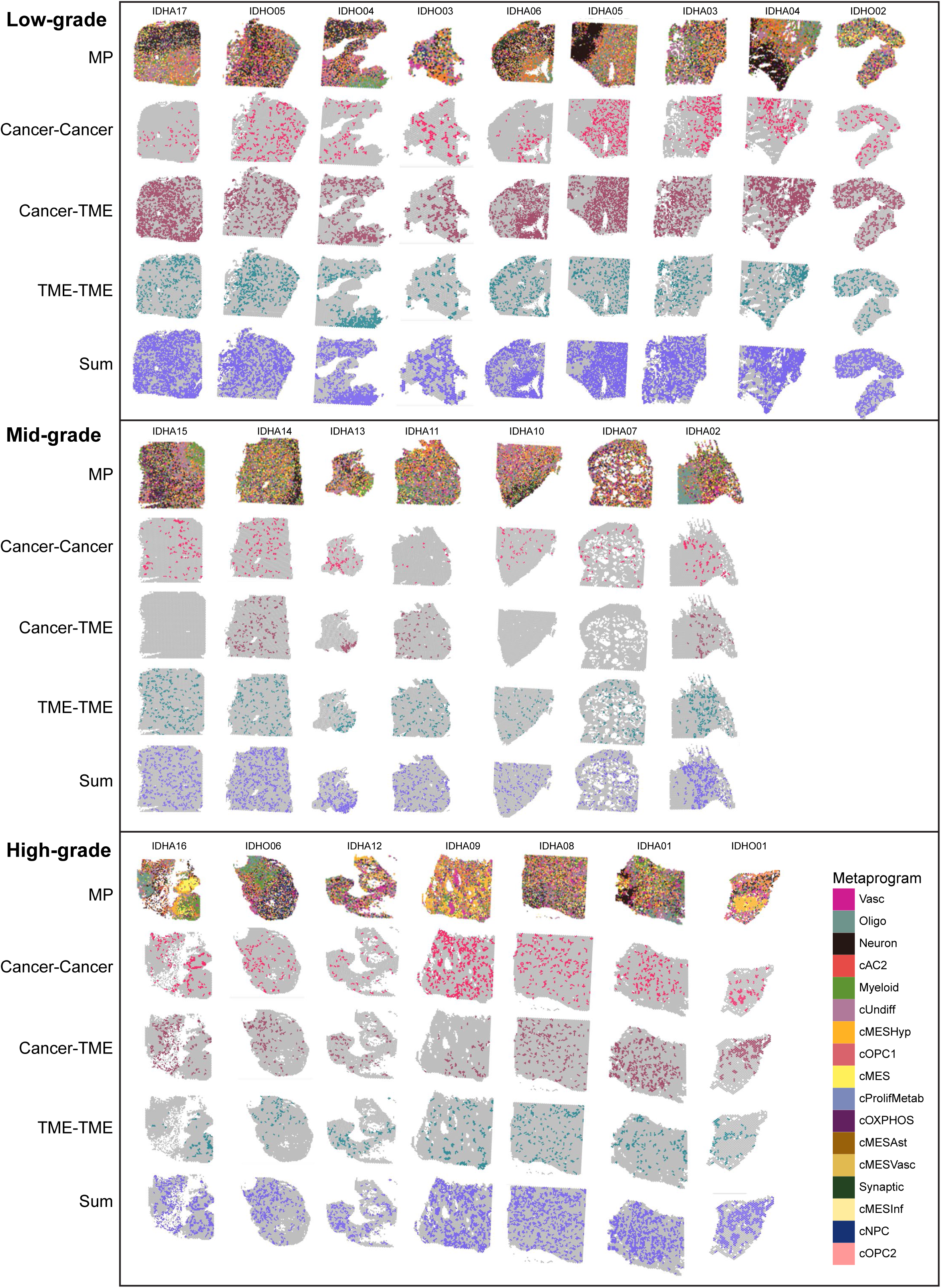
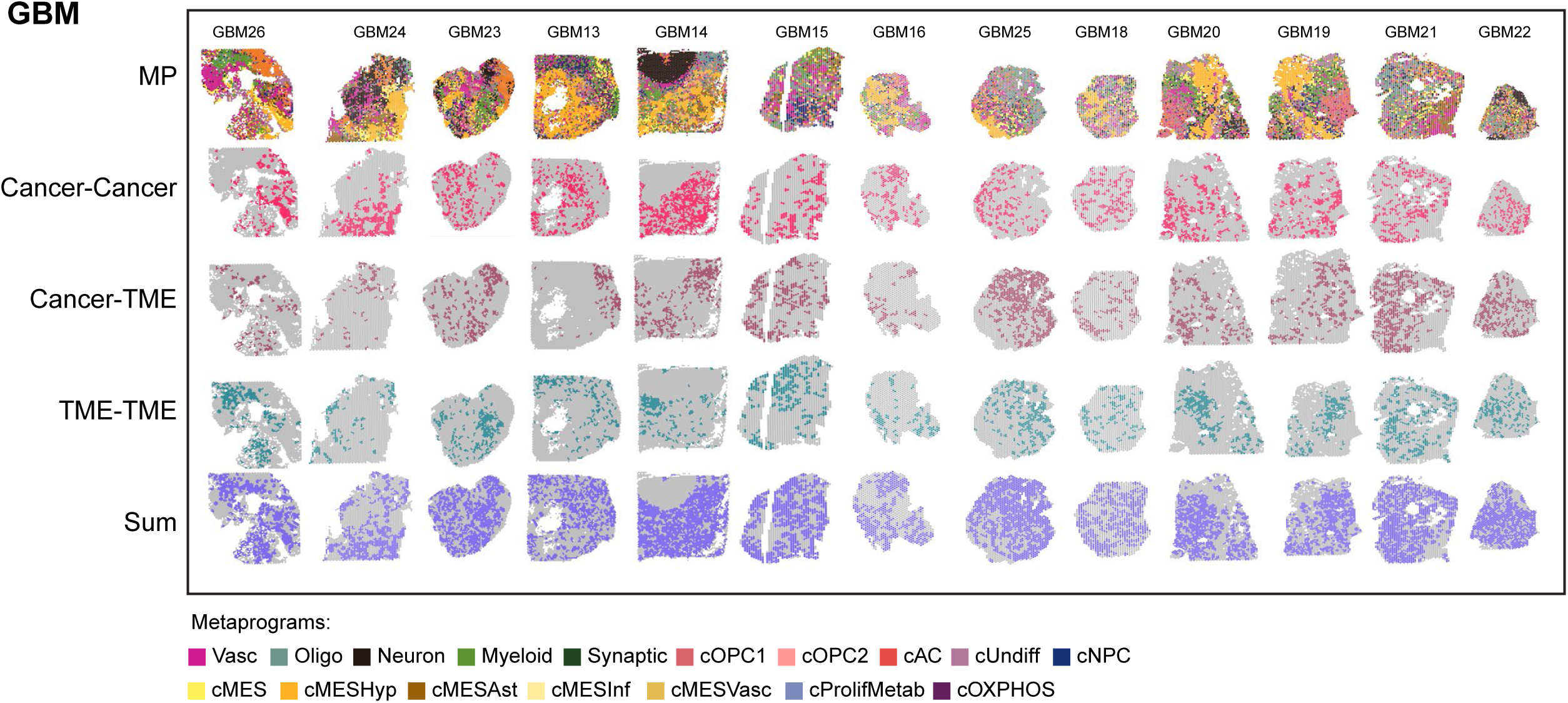
Visium spatial maps for al samples per categorical grade with five annotations: (i) Cell states (MPs); (ii) Total organziation; (iii) TME–TME consensus interactions; (iv) Cancer–TME consensus interactions; and (v) Cancer–Cancer consensus interactions.

## Supplemental Tables

**Table S1:** Patient and sample metadata, related to Figure 1.

**Table S2**: Gene signatures for Visium-derived metaprograms, related to Figure 1.

**Table S3:** CODEX panel and antibody specifications, related to Figure 2

## Methods

### Experimental Methods

#### Experimental model and study participant details

Glioma samples for spatial was collected from patients undergoing primary glioma resection at the Hospital St. Gallen, Switzerland, Massachusetts General Hospital, Boston, MA, Brigham and Women’s Hospital, Boston, MA, and at University Clinic Freiburg, Freiburg im Breisgau, Germany. All specimens were obtained with written informed consent and in accordance with institutional ethical approvals (KEK-ZH-Nr. 2015-0163; University Hospital Zurich; IRB #10-417; Dana-Farber Cancer Institute; University Hospital Freiburg IRB# 24-1100-S1; IRB #1360-1 at the Weizmann Institute of Science). Clinical characteristics of the cohort are summarized in Table S1.

#### Visium sample preparation

Fresh tumor samples were embedded in OCT at the time of freezing (Scigen OCT Compound, #4586). RNA integrity from multiple cryosectioned regions was assessed by TapeStation (Tapestation RNA ScreenTape, Agilent) following extraction with the Zymo Quick RNA MicroPrep Kit (#ZR-R1051). Only samples with RIN >7 proceeded to spatial transcriptomics profiling. Frozen tissue blocks were cryosectioned at 10 µm and placed onto the capture areas of 10x Genomics Visium Gene Expression slides following the manufacturer’s instructions (Slide & Reagent Kit PN-1000184). Sections mounted on Visium slides were stored at −80 °C and processed within two weeks.

#### Visium H&E Staining and imaging

Tissue sections on Visium slides were fixed in methanol (Millipore Sigma #34860), followed by an aqueous eosin-based H&E protocol according to manufacturer’s instructions (10X Visium Methanol Fixation, H&E Staining, and Imaging Protocol CG000160). Brightfield images were acquired on a Leica DMIi8 inverted microscope equipped with a DFC310FX color camera using a 10×/0.25 NA dry objective. Images were stitched using Leica Application Suite X and further processed in Fiji (v2.3.1).

#### Visium cDNA synthesis and library generation and sequencing

After imaging, tissue permeabilization was performed directly on the slide to release mRNA. Optimal permeabilization (9 minutes) was established using a Visium tissue-optimization time course. cDNA synthesis and library generation were performed with the Visium Spatial Gene Expression Slide and Reagent Kit (10X Genomics). Reverse transcription was performed by a template switch oligo protocol to generate a second strand, and cDNA synthesis was carried out according to qPCR results (KAPA FAST SYBR qPCR master mix, KAPA Biosystems). Amplified cDNA was fragmented, end-repaired, adaptor-ligated, and size-selected using SPRIselect beads (Beckman Coulter) at each step. Library quality was assessed using the Agilent TapeStation and NEBNext Library Quant Kit for Illumina (New England Biolabs). Indexed libraries were diluted to 1.8 nM, pooled, denatured, and sequenced on an Illumina NovaSeq SP flow cell (100-cycle kit) with 1% PhiX spike-in. Run configuration: Read 1 = 28 cycles; Read 2 = 90 cycles; Index 1 = 10 cycles; Index 2 = 10 cycles.

#### PhenoCyler Fusion

The antibody panel was curated to identify major cancer cell states and non-malignant cell types of the TME. We added markers to our previously published panel^9^ to increase phenotypic granularity in the myeloid compartment and the extracellular matrix. A detailed summary of all antibodies can be found in Table S3. Every antibody has been validated with conventional immunofluorescence staining using adequate control tissues from human duodenum and glioma, of which adjacent Visium data was available. PhenoCycler Fusion 2.0 (formerly CODEX; Akoya Biosciences) runs have been performed following the manufacturer’s instructions. In short, fresh frozen sections were fixed with 4% PFA for 20 min and blocked with 4% BSA and 0.25% TritonX-100 for 30 min. Primary antibodies were incubated overnight at 4 °C and secondary antibodies incubated for 2h at room temperature. All antibodies (including pre-conjugated antibodies from Akoya) were additionally validated on the respective glioma control samples using multiple CODEX runs to adjust antibody concentration and exposure times to optimize signal to noise ratio. After the run, hematoxylin and eosin staining was performed after flow cell removal with Xylene.

### Quantification and statistical analysis

#### Spatial transcriptomic analysis

##### Alignment, processing, and QC of Visium data

FASTQ alignment to GRCh38, UMI counting, and initial spot filtering were performed with Space Ranger (versions v1.2–v2.0, 10x Genomics). Spot-to-image alignment and spatial coordinate registration were carried out in Loupe Browser (v5.0.1).

Gene expression values were defined as *E_i_*_,*j*_ = log_2_ (1 + CPM*_i_*_,*j*_/10), in which counts per million (CPM)*_i_*_,*j*_ refers to 10^6^ × UMI*_i_*_,*j*_/sum(UMI_1…*n*,*j*_), for gene *i* in sample *j*, with *n* being the total number of analyzed genes. The average number of UMIs detected per spot was less than 100,000; thus, CPM values were divided by 10 to avoid inflating the differences between detected (*E_i_*_,*j*_ > 0) and undetected (*E_i_*_,*j*_ = 0) genes as previously described (Tirosh, 2016)

For each spot, the number of counts was used as a proxy for sample quality. Spots with fewer than 1000 counts and/or expressing more than 20% mitochondrial genes, another proxy for low quality, were filtered out. For each sample, the 7,000 most highly expressed genes were retained, and expression was centered per sample to define relative expression values by subtracting the average expression of each gene *i* across all *k* spots: Er*_i_*_,*j*_ = *E_i_*_,*j*_ − average(*E_i_*_,1…*k*_), where *Er* represents relative expression values.

##### Per sample clustering

Unsupervised clustering of spots was performed on each sample separately using two complementary approaches: Leiden clustering to capture discrete patterns of variation and NMF to capture continuous patterns of variation. For each sample, after PCA, Leiden clustering was performed on the SNN graph (Seurat v4.3.0). For Leiden clustering, cluster-specific gene programs were defined by differential expression analysis based on the top 50 most differentially expressed genes by the Wilcoxon Rank Sum Test with a *p*-value <0.005.

For NMF, negative values were transformed to zero in each centered expression matrix. To minimize the influence of selecting an individual *k* parameter, we performed NMF over multiple *k* values (2-10), generating 54 NMF programs per sample. Each NMF program was defined by the top 50 genes based on NMF score.

##### Generating consensus metaprograms

Gene programs from all samples (Leiden and NMF) were clustered based on their Jaccard overlap to identify consensus metaprograms (MPs). MPs were first defined for IDH-mutant gliomas alone, and a second set was subsequently defined that included both IDH-mutant and IDH-wild-type glioblastomas (including datasets from^9,11^) to enable cross-tumor-type comparisons.

Due to the high number of gene programs generated by NMF, programs were included in the consensus clustering only if they were (i) robust within a sample (i.e., 20/50 gene overlap between at least two programs across more than one value of k), (ii) non-redundant (i.e., for a group of similar programs, defined as at least 10% overlap with another program within a sample, only one is retained), and (iii) shared ≥10% gene overlap with programs from another sample. Clustering followed the procedures described in Gavish et al.^28^ and Greenwald et al.^9^.

##### Gene-set enrichment analysis

Visium-derived MPs generally map to a single cell type or cell state (see Greenwald et al. 2024 for validation of this approach). We assessed the enrichment of MP signatures with <u>Gene Ontology</u> terms (MSigDB modules H, C2, C5, C8) as well as published signatures for <u>glioma cell</u> states^3,9,12,29–31^, and non-malignant brain cell types^32–38^ by hypergeometric test (FDR adjusted *p* < 0.01 was considered significant).

##### Spot scoring and assignment of spots to MPs

Spot-wise MP scores were computed using the scalop R package (https://github.com/jlaffy/scalop). Given a set of genes (*Gj*) reflecting an MP corresponding to a cell state or cell type, we calculate for each spot *i*, a score, *SCj(i),* quantifying the relative expression of *Gj* in spot *i*, as the average relative expression (*Er*) of the genes in *Gj*, compared to the average relative expression of a control gene set (*Gj <u>cont</u>*): *SCj (i*) = average[*Er(Gj,i)*] – average[*Er*(*Gj cont,i*)]. The control gene-set is defined by first binning all analyzed genes into 20 bins of aggregate expression levels (*Ea*) and then, for each gene in the gene set *Gj*, randomly selecting 75 genes from the same expression bin. The control gene set has a comparable distribution of expression levels to that of *Gj*, and the control gene set is 100-fold larger, such that its average expression is analogous to averaging over 100 randomly selected gene sets of the same size as the considered gene set. Each spot was assigned to the MP with the highest score.

After quantifying malignant state composition per sample based on spot-based metaprogram annotations, we derived a per sample stemness score. For each sample, we calculated the proportion of each malignant state within the malignant compartment. The “stemness” score is defined as the proportion of “cUndiff”.

##### State specific coherence score

State-Coh was quantified independently for each metaprogram (MP) within each sample. For a given MP, we computed the mean number of immediately adjacent spots that shared the same MP assignment, averaged across all spots assigned to that MP. This value reflects the local self-association of the MP in space. To normalize this metric, we estimated a lower and upper bound on spatial coherence. The lower bound (expected minimum) was obtained by randomly permuting spot positions 100 times per sample and recalculating the mean same-MP neighbor count for each permutation; the shuffled results were then averaged to yield the expected value under spatial randomness. The upper bound (expected maximum) was defined analytically by assuming that all spots assigned to the MP were organized in a regular hexagon pattern, and computing the corresponding average number of same-MP neighbors under this configuration. The final spatial coherence score was computed by linearly scaling the observed value between these bounds:

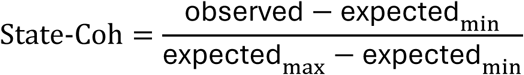

This normalization yields a score that facilitates comparison of State-Coh across MPs and samples.

##### Spot colocalization

For each sample, we first performed scoring-based spot deconvolution by assigning metaprogram (MP) scores to all spatial spots as described in *Spot scoring and assignment of spots to MPs*. MP scores above 0.1 were interpreted as indicating MP presence, and deconvolution matrices were binarized such that each MP was encoded as present (1) or absent (0) per spot. For any given MP pair (a, b) within a sample, we calculated the number of spots in which both MPs were present (N), as well as the total number of spots containing MP a (Ta) or MP b (Tb). Colocalization was quantified as:

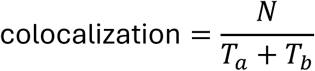

This observed value was averaged across spots to obtain a per-sample colocalization score for each MP pair. To estimate the expected colocalization, we repeated the same calculation on 500 shuffled versions of the deconvolution matrix per sample. Statistical significance was assessed using Fisher’s exact test by comparing observed and shuffled co-occurrence frequencies. An effect size was defined for each MP pair as the ratio of expected to observed colocalization. MP pairs were considered significantly colocalized within a sample if the effect size was ≥1.3 and the associated p-value was ≤0.01. Robustness across samples was evaluated by computing the fraction of samples in which each MP pair met these criteria. For downstream analyses, we used both the mean colocalization score per MP pair and the proportion of samples exhibiting significant colocalization.

##### Spot adjacency

Spatial adjacency was calculated independently for each sample. For a given ordered MP pair (A->B), we defined the observed asymmetric adjacency as the fraction of immediate neighbors of MP A spots that were assigned to MP B, normalized by all immediate neighbors of MP A:

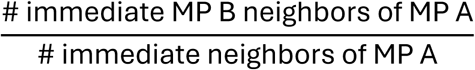

To remove the spatial coherence effect of MP B on its adjacency, we normalized this value by the relative adjacency capacity of MP B. Specifically, we calculated the proportion of MP B spots that had at least one neighboring spot not assigned to MP B, divided by the total number of such non-self adjacency opportunities across all MPs except MP A. To assess statistical significance, we generated a null distribution by randomly permuting spot positions 10,000 times per sample and recalculating adjacency scores. The *p*-value is determined by how many times the *observed* score was higher or lower than the shuffled scores (the smaller of the two).

##### Regional composition

Regional composition was calculated for each sample. For each radius (defined as the number of hexagons encircling a spot) ranging from 1 to 15, we defined a window around each spot. We then measured the abundance of all MPs within the window and computed correlations in abundance between MP pairs across all windows of the same size, excluding windows containing only one MP. This process was repeated over 500 shuffled permutations of the spot positions to generate a random distribution, from which we assessed the significance of the correlation using Fisher’s exact test.

##### Consensus interactions

To integrate spatial relationships across multiple scales, we rescaled the outputs of each spatial analysis (colocalization, adjacency, and regional composition) using min–max normalization, mapping all scores to a common range of −1 to 1. For each analysis, MP pairs with a mean scaled score greater than 0.35 across samples and that were statistically significant in at least 20% of samples within a given grade or tumor type were classified as *strongly connected*. An MP pair was defined as a ***consensus interaction*** if it satisfied the strong-connection criterion in at least two of the five spatial analyses (colocalization, adjacency, and regional composition at radii r = 5, 8, and 15), with the additional requirement that at least one measure reflected direct spatial coupling (colocalization or adjacency). MP pairs exhibiting consistent spatial avoidance were identified using the same criteria and classified as *repelled* interactions.

In order to validate that the overall low number of consensus interactions in mid-grade tumors is robust and to control for the differing number of samples per grade, we performed resampling and permutation controls for robustness using the proximity interaction statistics as computed from the spot adjacency measure per sample. To test whether differences in the number of recurrent interactions across grades could be explained by unequal group sizes, we down sampled each categorical grade to a common sample size (n=6) and repeated the procedure for six independent trials (random subsampling without replacement in each grade). For each down sampled group and trial, we computed the number of consistent enriched interactions as the number of metaprogram pairs with enrichment observed in at least three samples within the subsampled set and we computed the number of positive consensus-like interactions per trial. As a label-agnostic control, we generated six trials of random 4-way partitions. In each trial, we randomly selected six samples from each grade, permuted the pool samples, and split them into four groups of six samples each. For each group we computed proximity interactions and quantified the number of recurrent interactions. This analysis estimates the expected variability in recurrent interactions under random grouping when sample size and grade composition are controlled

##### Interaction category pairs

Consensus interactions and repulsions were further categorized by their interaction type (i.e., TME-TME; cancer-TME; and cancer-cancer) depending on the annotation of each MP as cancer (cOPC1, cOPC2, cAC, cUndiff, cNPC, cOXPHOS, cProlifMetab, cMESInf, cMESVasc, cMES, cMESAst, cMESHyp) or TME (Oligo, Neuron, Synaptic, Myeloid, Vasc). In order to compute the statistical significance of interaction category types within samples by grade and tumor type, we first quantified the relative abundance of each metaprogram pair participating in a consensus interaction across any grade or type (i.e., union of consensus interactions – global consensus pair) per sample. In order to quantify the relative abundance of each global consensus pair, we performed per-spot annotation of consensus interactions by checking whether the neighbor edges of each spot generated a consensus interaction with that spot. These per-spot annotations were stored both by metaprogram pair identity and by three category-level labels: Cancer-Cancer, Cancer-TME, and TME-TME. To test whether global consensus pair prevalence differs across groups and to generate type- and grade-specific consensus interaction annotations independent of our original definition of consensus interactions, we performed a Welch’s paired t-test on per-sample prevalence values by tumor grade and type. Grade comparisons included low/med/high, while type comparisons included IDH-A, IDH-O, and IDH-WT (GBM). We identified pairs whose prevalence differed significantly across grades or tumor types (p < 0.05). For pairs that were not significantly associated with a group, we retained them globally in all samples, and for pairs significantly associated with a given group, we retained them only for that group (collectively referred to as ‘allowed pairs’).

Allowed pairs were divided into interaction categories (Cancer-Cancer, Cancer-TME, TME-TME) using the pair-family annotation described above, and the relative abundance of each consensus interaction category among allowed pairs was calculated per sample. To test for differences in interaction composition across groups, we performed pair-wise Wilcoxon rank-sum tests on per-sample proportions (with Benjamini-Hochberg correction within each set of comparisons) and adjusted p-values<0.05 were considered significant.

#### Spatial proteomics analysis

##### Segmentation of spatial proteomics data

Raw images generated by the CODEX run were background subtracted as per the automated experimental post-processing. Pyramidal images were then loaded into QuPath^39^ and DAPI based segmentation with nuclear expansion by 2um was performed using CellPose^40^. In addition, an independent CellPose segmentation strategy based on DAPI and Vimentin was applied for the segmentation of gemistocytes. Mean expression values per segmentation mask were exported.

##### Nimbus scoring of spatial proteomics data

Next, multiplex images and segmentation masks were transformed into appropriate formats to be inputted into the Nimbus workflow^13^. Nimbus is a deep learning–based classifier trained on the Pan-Multiplex dataset, which scores marker positivity from multiplex imaging data. This approach avoids arbitrary intensity thresholds by leveraging pixel-level features to optimize positive/negative calls and is specifically suited to control for signal spillover of adjacent cells. We calculated Nimbus scores for every cell and every marker following the standard workflow.

##### Cell phenotyping of spatial proteomics data

Cell phenotyping was done in a two-step approach. First Nimbus scores of the most discriminatory markers were used to assign major cell types using a supervised and conservative gating strategy. Discriminatory markers were selected and ranked based on signal-to-noise ratio with an emphasis on nuclear markers as they are associated with less spillover artifact. We used the following gating strategy: Cells expressing OLIG2 > 0.4 were classified as cOPC. Oligo were defined by SOX10 > 0.6 and ATRX > 0.2 in the absence of OLIG2 expression (< 0.4). Neurons were defined by NeuN > 0.6. T cells were subdivided into CD8⁺ (CD3 > 0.7 and CD8 > 0.6) and CD4⁺ (CD3 > 0.7 and CD4 > 0.6) subsets., astrocytes by SPARC > 0.6 or S100B > 0.5 with ATRX > 0.7, and vascular cells by CD31 > 0.6 or HSPG2 > 0.5. Myeloid cells were identified by high expression of at least one of the following markers CD11c, CD163, CD206, or P2Y12 (> 0.6). Neutrophils and B cells were identified by CD45 > 0.7 co-expression with CD15 > 0.7 or CD19 > 0.7, respectively.

In a second step, we applied an unsupervised clustering workflow to the remaining unassigned cells. We used self-organizing maps (SOMs)^41^ using mean nuclear intensity values of representative state-specific marker genes (e.g., SOX2, ATRX, GFAP, VIM, S100B, CA9, PDPN). The resulting protein–cell matrices were transposed and used as input for SOM training on a two-dimensional rectangular grid (typically 3×3 or 3×2 units). Training was performed using Euclidean distance as the similarity metric over 10 iterations with learning rates decreasing from 0.05 to 0.01. Each cell was mapped to its closest SOM unit, effectively defining proteomically coherent subclusters. Cluster-level mean expression values and differential expression profiles were computed relative to all other clusters to identify the top marker genes for each SOM unit. Heatmaps of mean and z-scored differential expression were generated to visualize the expression patterns underlying cluster distinctions. To further assess intercluster relationships and subtype hierarchies, we performed hierarchical clustering of average expression profiles using Ward’s linkage. The resulting SOM-derived subclusters were then interpreted based on dominant marker gene expression patterns, and manually annotated as distinct proteomic states (e.g., cAC1, cAC2, Oligo, cUndiff) or filtered out as low-expression.

In a third step we used the same unsupervised SOM-based clustering approach to subtype T cells, Myeloid and Vasc using Nimbus scores of cell type specific markers, resulting in the final cell subtype annotations (**Fig. 2A**).

##### Colocalization analysis

We constructed neighborhood matrices for each sample using the imcRtools package in R. Neighborhoods were defined by a radius of 27.5 µm (corresponding to the diameter of one Visium spot). For each cell, the neighborhood matrix contained the counts of all neighboring cell types, with rows representing individual cells and columns the cell-type composition of their local neighborhood. Interaction frequencies between every possible pair of cell types were then summarized by normalizing counts to the abundance of the reference cell type within a sample.

To assess significance, we performed label-shuffling in which cell-type identities were permuted 500 times. Null distributions of interaction counts were derived from the shuffled data, and observed values were converted into z-scores relative to the mean and standard deviation of the null. Empirical *p*-values were calculated by comparing observed interaction strengths to the shuffled distributions, and significant cell–cell relationships were classified as either interaction or repulsion. Pairwise interaction profiles were then aggregated across samples and conditions to identify recurrent and condition-specific spatial relationships.

##### Junction analysis

###### TmnApp

We developed *TmnApp*, a Shiny/Leaflet tool for interactive exploration and annotation of single-cell or spot spatial maps. Users can filter views by glioma type, grade, and sample; color cells by annotation of cell types and states of various granularity or by e.g. IvyGAP; and optionally overlay marker expression (0–1 scaled). Regions of interest (ROIs) can be drawn directly on the map (polygon/line/circle) with two extend modes: (i) Extend Polyline, which generates parallel bands at user-defined thickness to sample along a line; and (ii) Extend Circle, which creates concentric annuli. ROI assignment is saved into the data frame for the selected sample and can be exported as CSV.

###### White and grey matter classification

Tissue sections stained with the CODEX proteomic panel were segmented into 5 µm tiles in QuPath, exporting mean marker intensities per tile. Expression matrices were log2(x+1)-transformed, followed by median centering and scaling across samples. To remove empty or artifact regions, we applied an unsupervised clustering workflow using a Self-Organizing Map and hierarchical clustering based on normalized marker expressions. Marker intensities were then averaged into 50 µm tiles, retaining those containing ≥ 10 small tiles.

Bimodal thresholds for PLP1 and MAP2 expression were determined using k-means clustering (k=2), corresponding to oligodendrocyte-associated (white matter) and neuronal (grey matter) markers, respectively. White and grey matter scores were computed as deviations from these thresholds, and each tile was annotated as white, grey, or unclassified based on relative expression dominance. This classification revealed that white matter compromised ∼70% and grey matter ∼30% of the total tissue area. This was validated by using additional neuronal markers (NeuN, SNAP25) and confirmed by IvyGAP annotation.

###### Junction cell density and purity analysis

Within each distance bin, the bin area was estimated from the bounding box of cell centroids and a fixed bin thickness of 1mm: area(mm²) = (length_µm × thickness)/10⁶. Cell density was computed as counts per mm². Purity was defined as the fraction of malignant cells. Distances were binned into four 1-mm intervals spanning −2 to +2 mm (Bin1 to Bin4). Bin2–Bin3 was defined as mid-zone baseline. Density values were expressed as relative change to this mid-zone baseline (Δ% Cells per mm²), while purity values were expressed as absolute deviation from baseline (Δ Purity). Per-sample behavior was visualized by connecting the bin-wise estimates with straight lines. Sample-level summaries (mean ± s.e. across samples) were plotted at each bin and linked with a black LOESS curve. A vertical reference line at 0 mm and a dashed horizontal reference line equal to the mean of the central bins (Bin2 and Bin3) were added. Paired t-tests comparing Bin1 vs Bin4 were used to assess significance, complemented by a *Consensus* metric indicating whether >50% of samples changed in a consistent direction (Consensus: TRUE).

###### Junction cell type composition analysis

The same distance binning was applied. For each sample, cell-type fractions were calculated by dividing the number of cells of a given type by the total cells within that bin, and relative changes were visualized as a per sample line plot. Sample summaries were obtained by averaging cell-type fractions across samples within each bin (mean ± s.e.), and these bin-wise means were connected with a black LOESS curve. A dashed red horizontal reference equals the average of the central bins (Bin2 and Bin3) for each facet. For each sample and cell type we estimated a per-sample null rate from the central region as the fraction in Bin2+Bin3. We then tested Bin1 and Bin4 against this null using binomial tests on counts (successes = cell-type counts; trials = total cells in the bin). For each bin we recorded the direction (above/below null) and *p*-value. If > 50% of evaluable samples agree in direction, we label significance by the fraction of samples with *p* < 0.05 and matching that majority direction: ≥75% = “**”, >50% = “*”, otherwise “NS”.

###### Junction marker expression analysis

We quantified relative expression of markers within a specific cell state (e.g. cOPC). For every sample and distance bin, relative marker expression scores were calculated and visualized as connected lines. Sample summaries were derived by averaging across samples within each bin (mean ± s.e.), and these bin-wise means were linked using a LOESS curve to display the overall trend. A per-marker “null” level was defined as the average of Bin2 and Bin3 means. For each marker, we formed paired per-sample comparisons between the extreme bins (Bin1 and Bin4). Let Δ = Bin4 − Bin1 be the within-sample change in relative Nimbus mean score. We report a two-sided paired *t*-test comparing Bin1 vs Bin4. Tests are computed only when ≥3 paired samples are available; otherwise, statistics are reported as NA. In addition, for each sample and marker we compared Bin1 and Bin4 to the per-marker null (Bin2 and Bin3) using two-sample Welch *t*-tests. For each bin we recorded direction (above/below the null) and *p*-value. Marker-level “votes” summarize consistency across samples: if a majority direction exists, we label significance by the fraction of samples with *p* < 0.05 and matching that direction: ≥75% = “**”, > 50% = “*”, otherwise “NS”.

## Declaration of generative AI and AI-assisted technologies in the manuscript preparation process

During the preparation of this manuscript, the authors used ChatGPT to assist with code organization and to improve stylistic clarity of the text. After using this tool, the authors reviewed and edited the content as needed and take full responsibility for the content of the published article.

